# Design and Development of DNA Damage Chemical Inducers of Proximity (DD-CIP) for Targeted Cancer Therapy

**DOI:** 10.1101/2025.11.03.686423

**Authors:** Tian Qiu, Yeuan Ting Lee, Brendan G. Dwyer, Yi Jer Tan, Ting Chen, Bryan A. Romero, Yanlan Wang, Jiehui Deng, Tinghu Zhang, Gerald R. Crabtree, Stephen M. Hinshaw, Kwok-Kin Wong, Nathanael S. Gray

## Abstract

Many chemotherapies are effective against cancers that display high levels of genome instability by disrupting or overwhelming the DNA damage response to induce cell death. PARP inhibitors (PARPi) exploit this vulnerability by stalling DNA repair particularly in homologous recombination (HR)-deficient cancer cells. Although PARPi are now used to treat BRCA1/2-mutated cancers such as ovarian and breast cancers, they are still limited to a narrow range of clinical indications and are susceptible to acquired resistance. Here, we introduce “DNA Damage Chemical Inducers of Proximity” (DD-CIPs), bivalent molecules that rewire the mechanism of action of conventional PARPi. The DD-CIPs function through chemical induced proximity between PARP1/2 and the chromatin remodeling protein, BRD4. From a candidate library of DD-CIPs, we identified DD-CIP1 which induces the DNA damage response (DDR) and apoptosis to a range of cancer lines at two-digit nanomolar concentrations. Further optimization yielded DD-CIP2, which induces tumor cell death at nanomolar concentrations across diverse blood and solid cancer cells, including cancer types that are insensitive to PARPi. Using small-cell lung cancer (SCLC) as a model, we found that DD-CIP2 triggers DDR, cell cycle arrest, and apoptosis *in vitro,* leading to anti-tumor efficacy without substantial toxicity in preclinical SCLC xenograft models at well tolerated doses. Our findings demonstrate that DD-CIPs may provide an opportunity to address the limitations of traditional PARPi and establish chemical induced proximity as a strategy for modulating the DDR in cancer.

## Main

Cancers commonly harbor elevated levels of DNA damage relative to normal tissue^1^. Consequently, cancer cells frequently upregulate DNA repair pathways and repress normal cell-cycle checkpoints and apoptotic signaling^2^. These adaptations distinguish malignant cells from normal cells, presenting opportunities for therapeutic intervention. Conventional DNA-damaging agents such as cisplatin and doxorubicin exploit these vulnerabilities by inducing DNA lesions that are lethal to fast-dividing cancer cells^3^. Despite broad clinical use, these agents produce systemic toxicity that often limits dose escalation^4, 5^. Recently, poly(ADP-ribose) polymerase 1 (PARP1) inhibitors have emerged as targeted therapies that exploit homologous recombination (HR) deficiencies, particularly in BRCA1/2-mutant cancers, offering an alternative to conventional genotoxic chemotherapy^6, 7^. However, their clinical benefit has remained largely confined to patients with HR-deficient cancers, underscoring the need for new therapeutic modalities with broader efficacy and improved targetability.

Chemically induced proximity (CIP) offers a promising strategy for reprogramming protein-protein interactions (PPIs) with small molecules, enabling event-driven modulation of diverse cellular processes such as protein re-localization^8–10^, signal transduction^11^ and blockade^12–14^, transcriptional and epigenetic regulation^15–20^, and post-translational modification^21–27^. Our group recently expanded this modality by introducing transcriptional/epigenetic CIPs (TCIPs), which recruit the transcriptional regulators such as BRD4^17, 28^, CDK9/pTEF-B^29^, or p300^30^, to promoters bound by BCL6 or TP53, thereby restoring apoptotic or cell-cycle arrest transcriptional programs in cancer cells.

To overcome the limitations of conventional DNA-damaging agents, we explored whether CIPs could be engineered to rewire the DNA damage response (DDR) in cancer cells by recruiting cancer-relevant effector proteins directly to the sites of DNA damage. We focused on PARP1, a primary DNA damage sensor that detects both single- and double-stranded breaks and contributes to the repair cascades^6^. Amongst the clinically approved PARP inhibitors (PARPi), type II inhibitors trap PARP1 on DNA damage lesions by prolonging its DNA-bound state^31^, making them ideal ligands for anchoring CIPs to damaged DNA. Prior studies have shown that BRD4 inhibition is synthetic lethal with PARPi by suppressing HR^32, 33^, and dual PARP1/BRD4 inhibitors have been developed based on this synergy^34–37^, yet their activity in HR-proficient cancers remains similar to co-treatment with PARP1i and BRD4 inhibitors. Building on our prior work showing that TCIP1 rapidly recruits BRD4 to chromatin^17^, we tested whether directing BRD4 to PARP1 could generate a synthetic protein complex that interferes with DDR and thereby might enhance cancer cell killing. This strategy represents a mechanistically distinct approach to conventional PARPi or dual inhibitors and may establish a precedent for recruiting oncoproteins or chromatin modulators to PARPi in the future.

Here, we developed “DNA Damage Chemical Inducers of Proximity” (DD-CIPs) to accomplish this objective. Our lead compound, DD-CIP1, was generated by connecting a type II PARPi (Olaparib) to a BRD4-recruiting ligand (JQ1) via a flexible alkyl linker. In cancer cells, DD-CIP1 exhibited cytotoxicity comparable to or exceeding that of co-treatment with the parental PARP and BRD4 inhibitors. Mechanistic studies demonstrated that DD-CIP1 induces association between BRD4 and PARP1/2 and enhances DNA damage response and apoptosis in cancer cells. We further developed DD-CIP2 by introducing a rigidified linker, which displayed superior killing potency against multiple blood- and solid tumor-derived cancer cell lines regardless of their BRCA1/2 status. Using small cell lung cancer (SCLC) as model, we found that DD-CIP2 induces robust DNA damage and apoptosis *in vitro* and exhibits anti-tumor efficacy *in vivo*. These findings position DD-CIPs as a promising therapeutic modality that leverages proximity pharmacology to target cancers.

## Results

### Development of DD-CIP1

To assess the feasibility of targeting the DDR through induced proximity, we designed and synthesized a series of bivalent molecules that recruit BRD4 to PARP1 (**Figure 1A, S1**). These compounds were generated by covalently linking PARP1 inhibitors (Olaparib, Rucaparib, Veliparib or Niraparib) to the BRD4 inhibitor JQ1 via linkers of varying lengths and chemical compositions.

**Figure 1.**
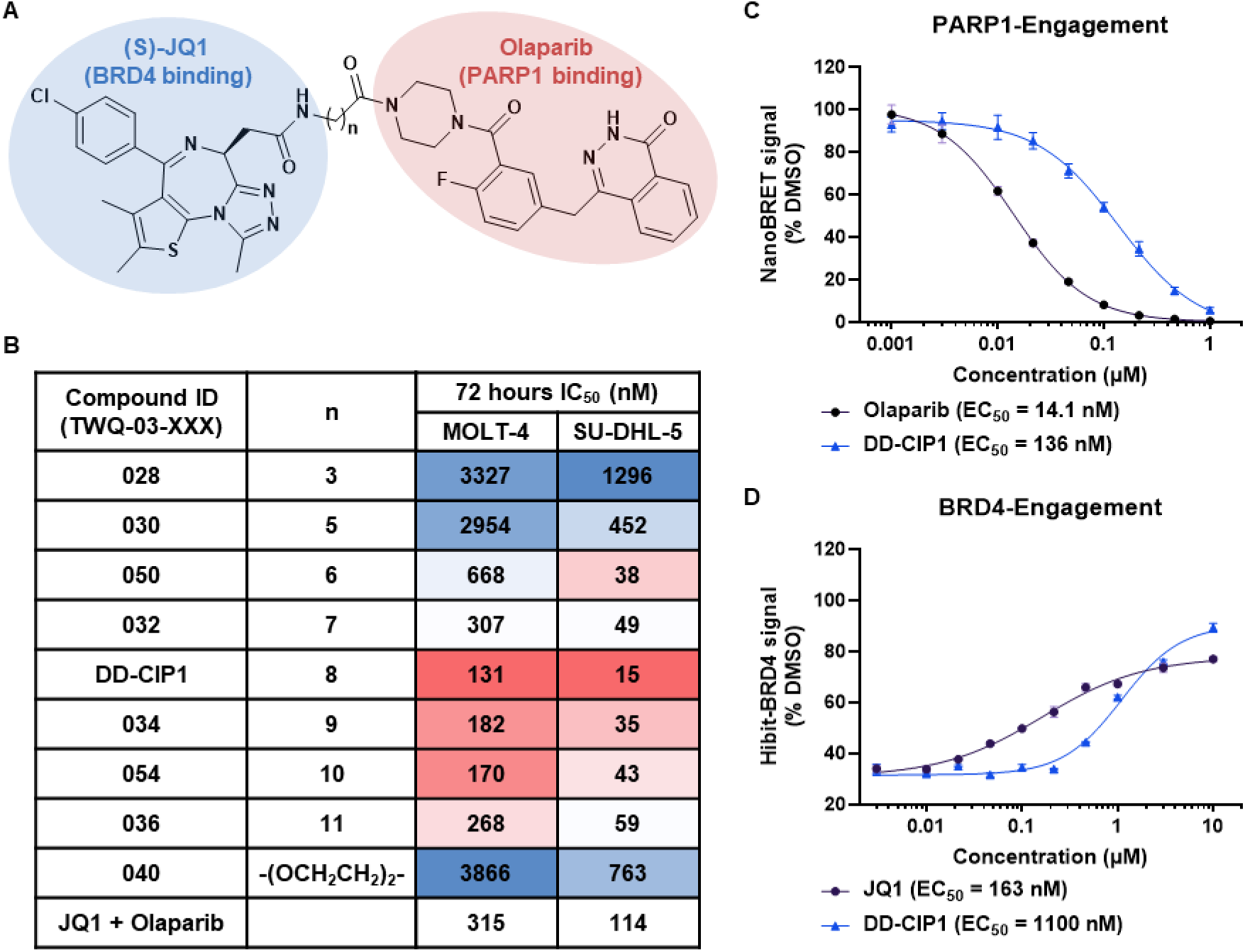
Development of DD-CIP1. A) Chemical structure of DD-CIPs. B) Cell-killing potencies of DD-CIP compounds compared to the co-treatment of JQ1 and Olaparib (1:1) at 72 hours in SU-DHL-5 and MOLT-4 cells. Data are shown as means; n = 3 biological replicates. C) PARP1 target engagement assay in HEK293T cells measured by Olaparib or DD-CIP1 competitive displacement of Tracer-PARP-01 (NanoBRET signal normalized to DMSO. Data are shown as means ± SEM; n=3 biological replicates). D) BRD4 target engagement assay in Hibit-BRD4 Jurkat cells measured by JQ1 or DD-CIP1 competitive displacement of dBET6 (HiBiT luminescence normalized to DMSO control with no dBET6; Data are shown as means ± SEM; n=3 biological replicates).

To identify the most potent compounds, we conducted a cell viability screen in HR-proficient blood cancer cell lines, MOLT-4 (T-cell acute lymphoblastic leukemia) and SU-DHL-5 (B-cell lymphoma), using a CellTiter-Glo assay. DD-CIP1, composed of Olaparib, JQ1 and an 8-carbon (C8) alkyl linker, emerged as the most potent molecule, with IC_50_ values of 131 nM and 15 nM for MOLT-4 and SU-DHL-5, respectively. These potencies surpassed the efficacy of co-treatment with the parental compounds (315 nM and 114 nM, respectively). Linker composition modulated cytotoxicity: optimal cytotoxic activity was observed with C8-C10 alkyl chains, and shorter or longer linkers reduced the activity (**Figure 1B**). Substitution of alkyl linker with PEG-based linker (TWQ-03-040) or modification of the Olaparib exit vector from an amide to an alkyl linkage (TWQ-03-086) led to a loss in potency (**Figure S1A**). This matches trends observed in PARP1-targeting PROTACs, where aliphatic linkers are generally favored^38, 39^. Additional analogs employing Rucaparib, Veliparib or Niraparib exhibited similar activity compared with co-treatment but were less potent than DD-CIP1 (**Figure S1B**). We also synthesized two negative control compounds containing the DD-CIP1 linker but with impaired binding to PARP1^40^ (TWQ-03-064) or BRD4^41^ (TWQ-03-065) (**Figure S2A**). Both control compounds showed reduced potency compared with DD-CIP1 (**Figure S2B**). Based on these findings, we selected DD-CIP1 for further characterization.

To investigate the mechanism of action of DD-CIP1, we used cellular target-engagement assays to measure the ability of DD-CIP1 to bind BRD4 or PARP1. In the PARP1 target engagement assay, DD-CIP1 was approximately 10-fold less effective than Olaparib in displacing a fluorescent tracer from PARP1-Nanoluc (**Figure 1C, S3A**). Likewise, in the BRD4 target engagement assay, DD-CIP1 was about 7-fold less effective at rescuing dBET6 induced BRD4 degradation as measured by the HiBiT-BRD4 assay^42^ (**Figure 1D, S3B**). These target engagement studies indicate that the improved anti-cancer activity of DD-CIP1 is not due to improved engagement with BRD4 or PARP1, eliminating the possibility of a RIPTAC-like mechanism^14^.

### DD-CIP1 induces PARP1-BRD4 binding

We hypothesized that DD-CIP1 cellular activity may derive from its ability to serve as a molecular glue between PARP1 and BRD4. Support for this idea came from our observation of strong linker length dependency in the initial structure-activity relationship studies. To test this, we used co-immunoprecipitation (co-IP) experiments in HEK293T cells transiently expressing FLAG-tagged BRD4. PARP1 pull-down using anti-FLAG M2 magnetic beads was only observed when cells were treated with DD-CIP compounds that contain more than 5-carbon (C5) linker (**Figure 2A**), which matches the linker-dependent efficacy profile observed in cell viability assays. Reciprocally, co-IP of FLAG-PARP1 in the presence of DD-CIP1 pulled down both the long and short isoforms of endogenous BRD4 (**Figure 2B**), while treatment with JQ1 did not lead to the same phenotype. Thus, DD-CIP1 could induce robust PARP1–BRD4 interaction.

**Figure 2.**
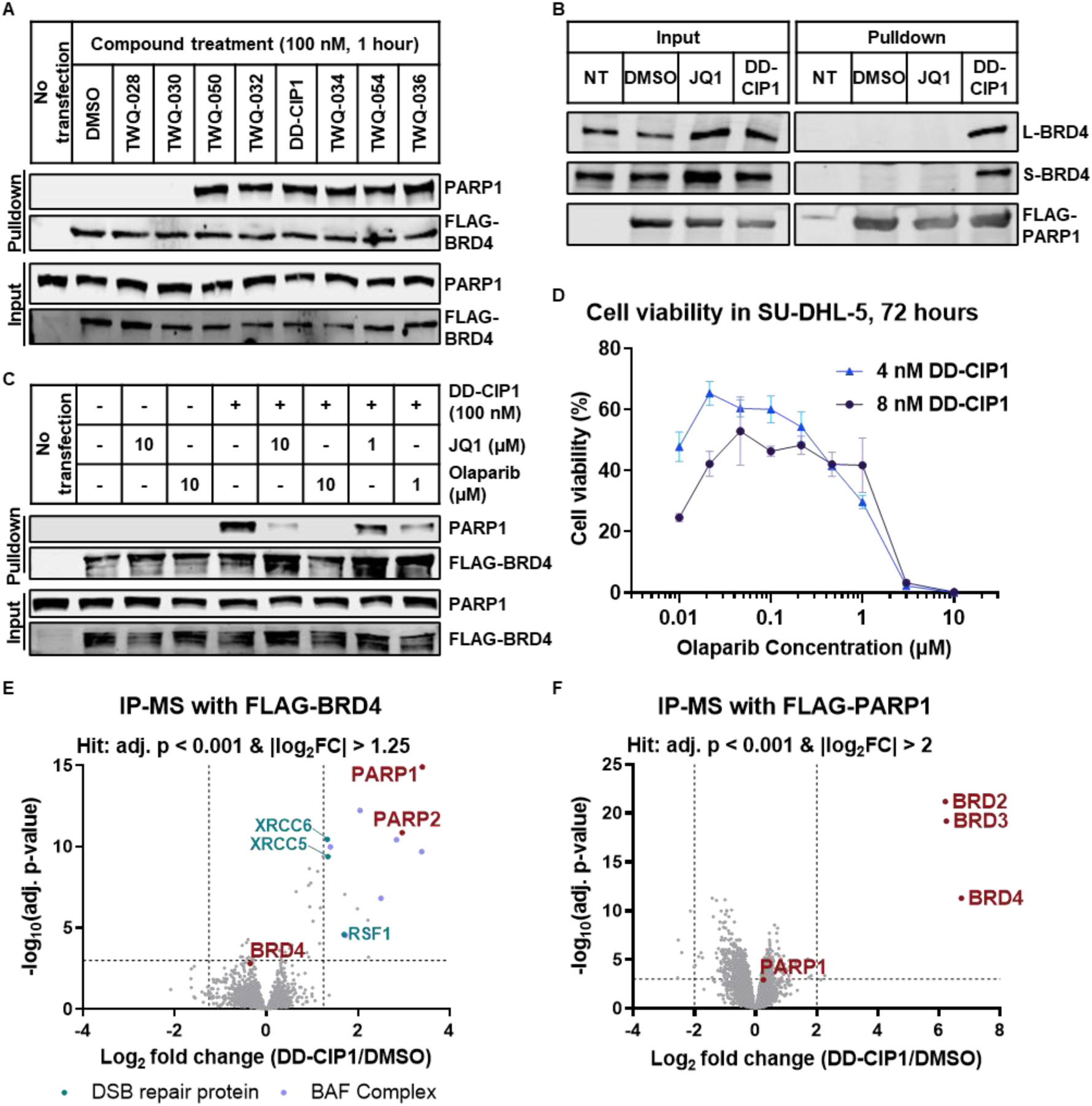
DD-CIP1 induces neo-interaction between PARP1 and BRD4. A) Co-immunoprecipitation (Co-IP) of FLAG-BRD4 and PARP1 in HEK293T cells after 1 hour treatment of 100 nM DD-CIPs. B) Co-IP of FLAG-PARP1 and BRD4 in HEK293T cells after 1 hour treatment of 1 μM JQ1 or DD-CIP1. NT indicates no transfection of FLAG-PARP1. L-BRD4 and S-BRD4 indicate long and short isoform of BRD4, respectively. C) Co-IP of FLAG-BRD4 and PARP1 in HEK293T cells. The cells were pretreated with JQ1 or Olaparib for 1 hour at indicated concentrations, followed by the treatment of 100 nM DD-CIP1 for 1 hour. D) Measurement of SU-DHL-5 cell viability after competitive titration of Olaparib to constant 4 or 8 nM DD-CIP1; cells were treated simultaneously with DD-CIP1 and Olaparib for 72 hours. Data are shown as means ± SEM; n = 3 biological replicates. E) FLAG Immunoprecipitation-mass spectrometry (IP-MS) in HEK293T cells overexpressing FLAG-BRD4 treated with 100 nM DD-CIP1 for 1 hour; plotted with cut-offs of |log_2_(fold change) | ≥ 1.25 and adj. P ≤ 0.001; 3 biological replicates. F) FLAG IP-MS in HEK293T cells overexpressing FLAG-PARP1 treated with 100 nM DD-CIP1 for 1 hour; plotted with cut-offs of |log_2_(fold change) | ≥ 2 and adj. P ≤ 0.001; 3 biological replicates. All P values were adjusted using Benjamini-Hochberg method from LIMMA-moderated t-test.

To more precisely characterize DD-CIP1 induced neo-interaction, we conducted time- and dose-dependent co-IP experiments in HEK293T cells overexpressing FLAG-BRD4. DD-CIP1 mediated PARP1-BRD4 association at concentrations below 10 nM, with maximal interaction formed at 50 nM after 1 hour of treatment (**Figure S4A**). A time-course experiment at 50 nM demonstrated a rapid complex assembly, and the equilibrium was reached after 40 minutes (**Figure S4B**). To confirm protein interactions in live cells, we used NanoBRET assay to measure the ternary complex formation between Nanoluc-BRD4 and HaloTag-PARP1 in HEK293T cells. DD-CIP1 treatment produced a dose-dependent NanoBRET signal, peaking at 100 nM and displaying a bell-shaped curve characteristic of the saturation behavior of bivalent molecules at high concentrations^43, 44^ (**Figure S4C**).

We next investigated whether the ternary complex is required for the cell killing properties of DD-CIP1. Pretreatment of cells with JQ1 or Olaparib decreased PARP1-BRD4 association in co-IP experiments, suggesting both sides are necessary for ternary complex formation (**Figure 2C**). Titrating increased concentrations of the PARPi Veliparib or Olaparib to SU-DHL-5 cells, against constant concentrations of DD-CIP1, attenuated the cytotoxicity of DD-CIP1, suggesting ternary complex-dependent cytotoxic activity (**Figure 2D, S4D**). Collectively, these data indicate that DD-CIP1 rapidly induces PARP1-BRD4 association which contributes to *in vitro* cytotoxicity.

The human genome encodes 17 PARP paralogs, and Olaparib targets several of them^45^. To determine which PARP protein is associated with DD-CIP1 activity in cells, we carried out immunoprecipitation-mass spectrometry (IP-MS) in HEK293T cells overexpressing FLAG-BRD4. Briefly, the cells were treated with 100 nM DD-CIP1 for 1 hour, lysed and the lysate were subjected to anti-FLAG M2 magnetic beads, followed by label-free quantitative proteomics to identify interactors. PARP1 and PARP2 were among the most enriched proteins (adj. p < 0.001, |log_2_FC| > 1.25; **Figure 2E; Supplemental Table 1**). We also identified co-enrichment of several DNA double strand break (DSB) related proteins, including XRCC5, XRCC6, SSBP1 and RSF1, which are commonly localized to DSB sites and can be modulated by PARP1^46, 47^. These data suggest that DD-CIP1 treatment may re-localize BRD4 to the DNA break sites. To further assess bromodomain protein engagement, we performed IP-MS in HEK293T cells overexpressing FLAG-PARP1. Here, BRD2, BRD3, and BRD4 were strongly enriched with DD-CIP1 treatment (adj. p < 0.001, |log_2_FC| > 2; **Figure 2F**), consistent with the known binding profile of JQ1 across bromodomain family members^41^. As a control, Olaparib treatment in HEK293T cells overexpressing FLAG-PARP1 did not result in significantly enriched proteins (**Figure S4E**).

### DD-CIP1 activates DNA damage response and induces apoptosis

Next, we monitored DNA damage response (DDR) by measuring γH2AX and apoptosis by PARP1 cleavage using Western blot. In SU-DHL-5 cells treated with DD-CIP1 for 24 hours, we observed increased γH2AX and cleaved PARP1, whereas cells co-treated with JQ1 and Olaparib showed much weaker response (**Figure 3A**). To further investigate whether DD-CIP1 accelerates DDR signaling, we treated SU-DHL-5 cells with 200 nM DD-CIP1 or the combination of JQ1 and Olaparib and harvested cells over a 15 min to 6 hours’ time period. DD-CIP1 led to the phosphorylation of ATM, ATR, CHK1 and CHK2, the commonly used DDR markers^48, 49^, as early as 30 min post treatment (**Figure 3B**). By contrast, JQ1 and Olaparib co-treatment produced minimal DDR activation within the same timeframe, consistent with that the traditional PARP inhibitors typically require prolonged exposure to drive DDR signaling^50^. These results suggest that DD-CIP1 can induce a rapid DDR in cells, which also aligns with the fast kinetics of BRD4 recruitment to PARP1.

**Figure 3.**
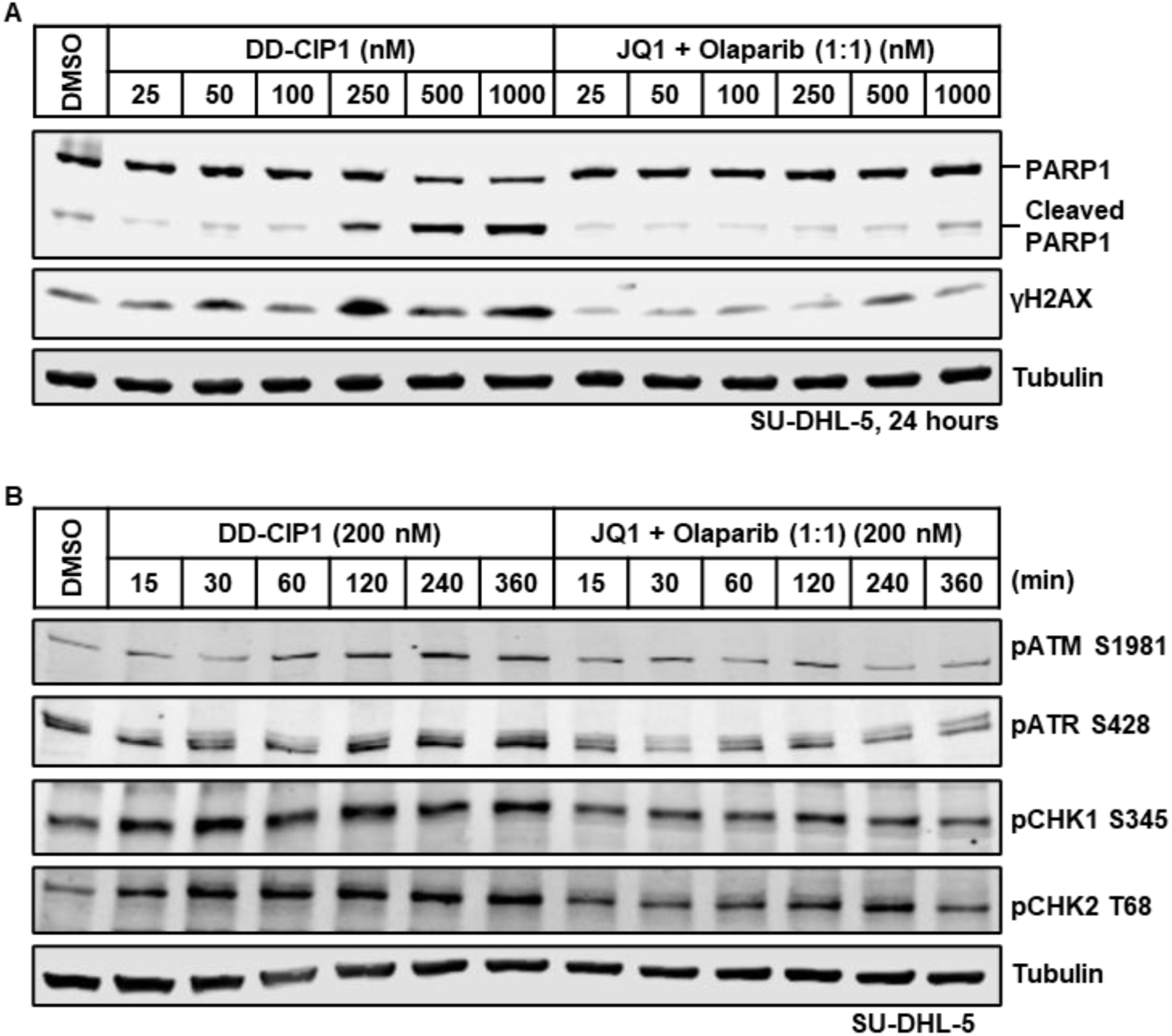
DD-CIP1 induces rapid DNA damage response and activates apoptotic signals. A) Western blots showing apoptosis (cleaved PARP1) and DNA damage (γH2AX) markers in SU-DHL-5 cells treated with the indicated concentration of DD-CIP1, or co-treatment of JQ1 and Olaparib (1:1) for 24 hours. B) Western blots showing phosphorylation of DNA damage response proteins in SU-DHL-5 cells treated with 200 nM of DD-CIP1, or 200 nM co-treatment of JQ1 and Olaparib for indicated time.

### DD-CIP2 with a rigidified linker exhibits improved potency and bio-stability

While DD-CIP1 demonstrated proof-of-concept for neo-interaction dependent cytotoxicity, its poor microsomal stability may limit its usefulness as a pharmacological probe (**Figure S5**). To address this, we prepared a small library of DD-CIP1 analogs with varied linkers and identified DD-CIP2, which features a glycine-conjugated 3,9-diazaspiro[5.5]undecyl linker (**Figure 4A**). DD-CIP2 demonstrated dramatically increased potency in SU-DHL-5 and MOLT-4 cells versus DD-CIP1 (EC_50_ = 0.78 nM and 4.6 nM, respectively; **Figure 4B**, **Table 1**), representing a ∼23-fold and ∼28-fold enhancement anti-proliferative activity relative to of DD-CIP1.

**Figure 4.**
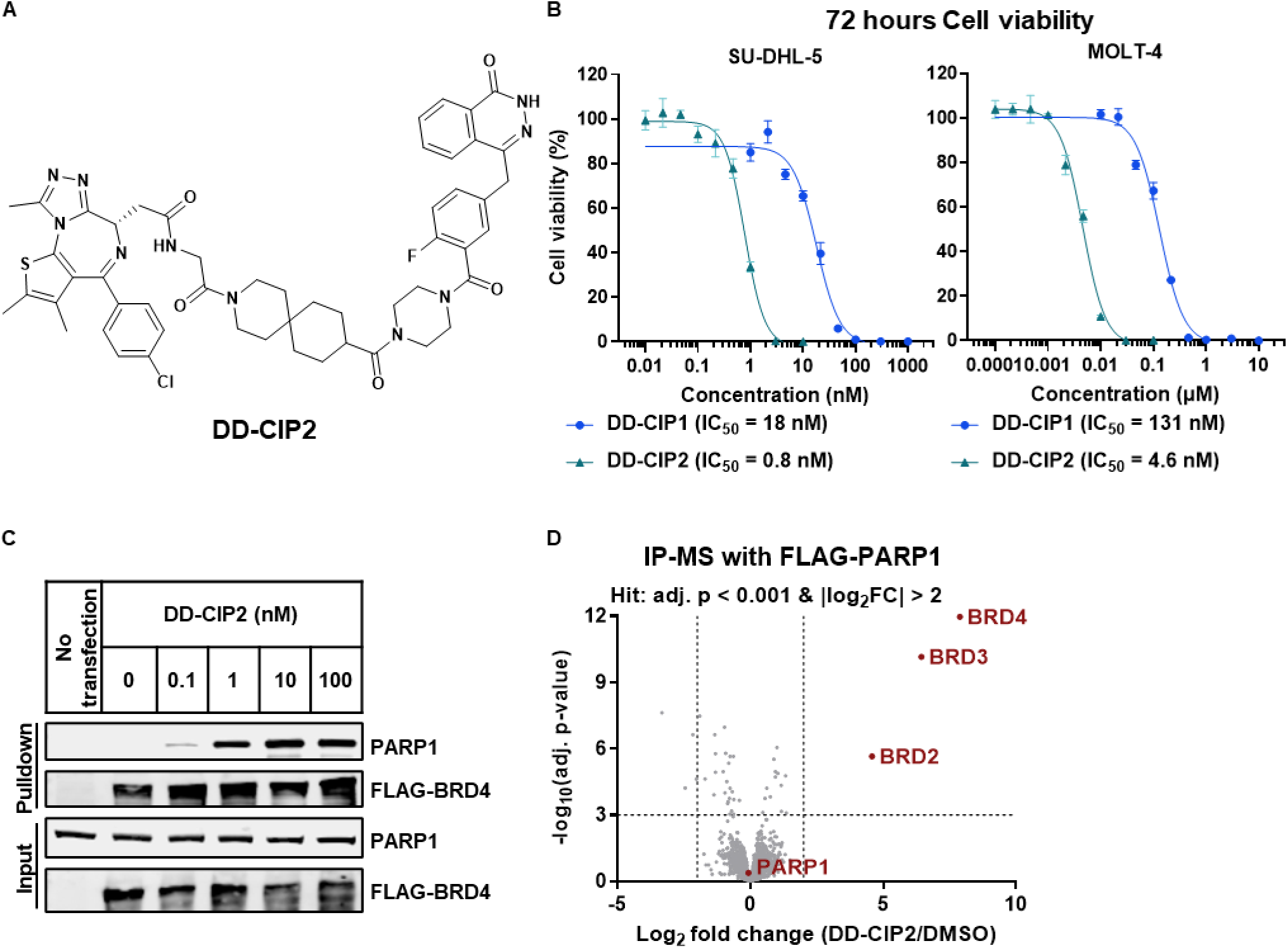
DD-CIP2 with a rigidified linker exhibits improved potency. A) Chemical structure of DD-CIP2. B) Cell-killing potencies of DD-CIP1 and DD-CIP2 after 72 hours treatment in SU-DHL-5 and MOLT-4 cells. Data are shown as means ± SEM; n = 3 biological replicates. C) Co-IP of FLAG-BRD4 and PARP1 in HEK293T cells after 1 hour treatment of indicated concentrations of DD-CIP2. D) FLAG IP-MS in HEK293T cells overexpressing FLAG-PARP1 treated for 1 hour with 100 nM DD-CIP2; plotted with cut-offs of |log_2_(fold change) | ≥ 2 and adj. P ≤ 0.001; 3 biological replicates. All P values were adjusted using Benjamini-Hochberg method from LIMMA-moderated t-test.

**Table 1.**
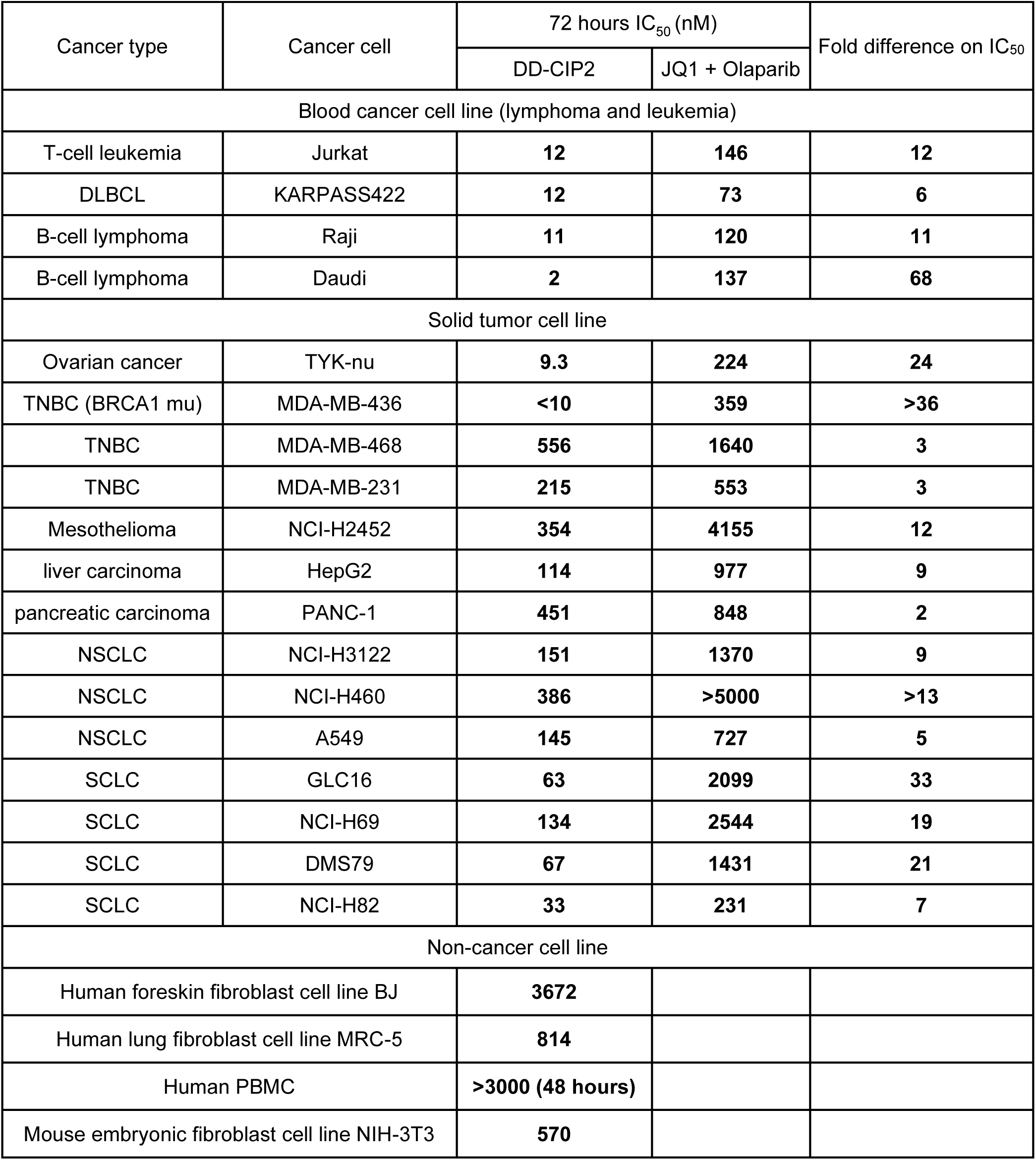
Efficacy of DD-CIP2 compared to co-treatment of JQ1 and Olaparib in different cell lines (Data are shown as means, n = 3)

Compared to DD-CIP1, DD-CIP2 displayed only a modest increase of PARP1 occupancy (∼3-fold) and similar BRD4 occupancy from cellular engagement assays (**Figure 1B,C, S6A,B**), demonstrating that the superior cytotoxic activity of DD-CIP2 was not solely attributable to stronger target binding versus DD-CIP1. Like DD-CIP1, DD-CIP2 induced PARP1-BRD4 interaction, as confirmed by IP-Western blot (**Figure 4C**). Co-treatment with the Olaparib rescued the effects of DD-CIP2 on cell viability, demonstrating the dependency on the PARP-BRD4 association (**Figure S6C**). IP-MS experiments revealed similar protein interaction profiles for DD-CIP2 to DD-CIP1 (**Figure 4D, S6D**). Pharmacokinetic (PK) study showed that DD-CIP2 has a favorable half-life of 2.3 hours following intraperitoneal (IP) administration, supporting that it would be suitable as a pharmacological probe for use in murine tumor models (**Table 2**). Together, these findings suggest that DD-CIP2 is a potent and bioavailable small molecule that retains the DD-CIP1’s mechanism of action.

**Table 2.**
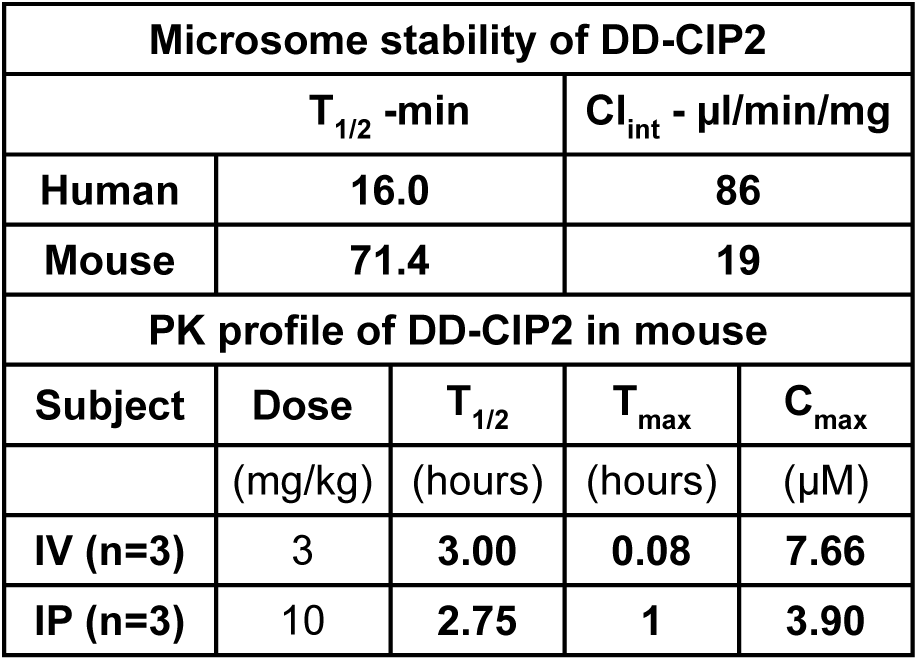
Microsome stability of DD-CIP2 and PK profile of DD-CIP2 in mouse.

We next investigated the therapeutic potential of DD-CIP2 by examining its activity in a diverse panel of blood cancer and solid tumor cell lines (**Table 1**). DD-CIP2 exhibited enhanced potency compared to JQ1 and Olaparib co-treatment, achieving over 10-fold greater efficacy in more than half of the tested cancer cell lines. DD-CIP2 was also active in cancer types that are generally insensitive to co-treatment with JQ1 and Olaparib, such as Mesothelioma, NSCLC and SCLC cells, highlighting its broad application. DD-CIP2 produced comparatively low cytotoxicity in untransformed human and mouse fibroblast cells and human PBMC (Table 1), supporting its potential for cancer targeting rather than general toxicity. Together, these findings suggest that DD-CIP2 can target a broad spectrum of cancers.

### DD-CIP2 effectively induces DNA damage response and anti-tumor efficacy in SCLC model in vitro and in vivo

To further evaluate the therapeutic potential of DD-CIP2, we selected SCLC as a representative solid tumor model because: (1) BRCA1/2 mutations are rare in SCLC^51^, providing an opportunity to evaluate DD-CIP2 activity in a homologous recombination (HR)-proficient context; and (2) DD-CIP2 demonstrated approximately 10-fold enhanced cytotoxicity compared with co-treatment of parental inhibitors in SCLC lines (GLC16, NCI-H69, DMS79, and NCI-H82; **Figure 5A-D**).

**Figure 5.**
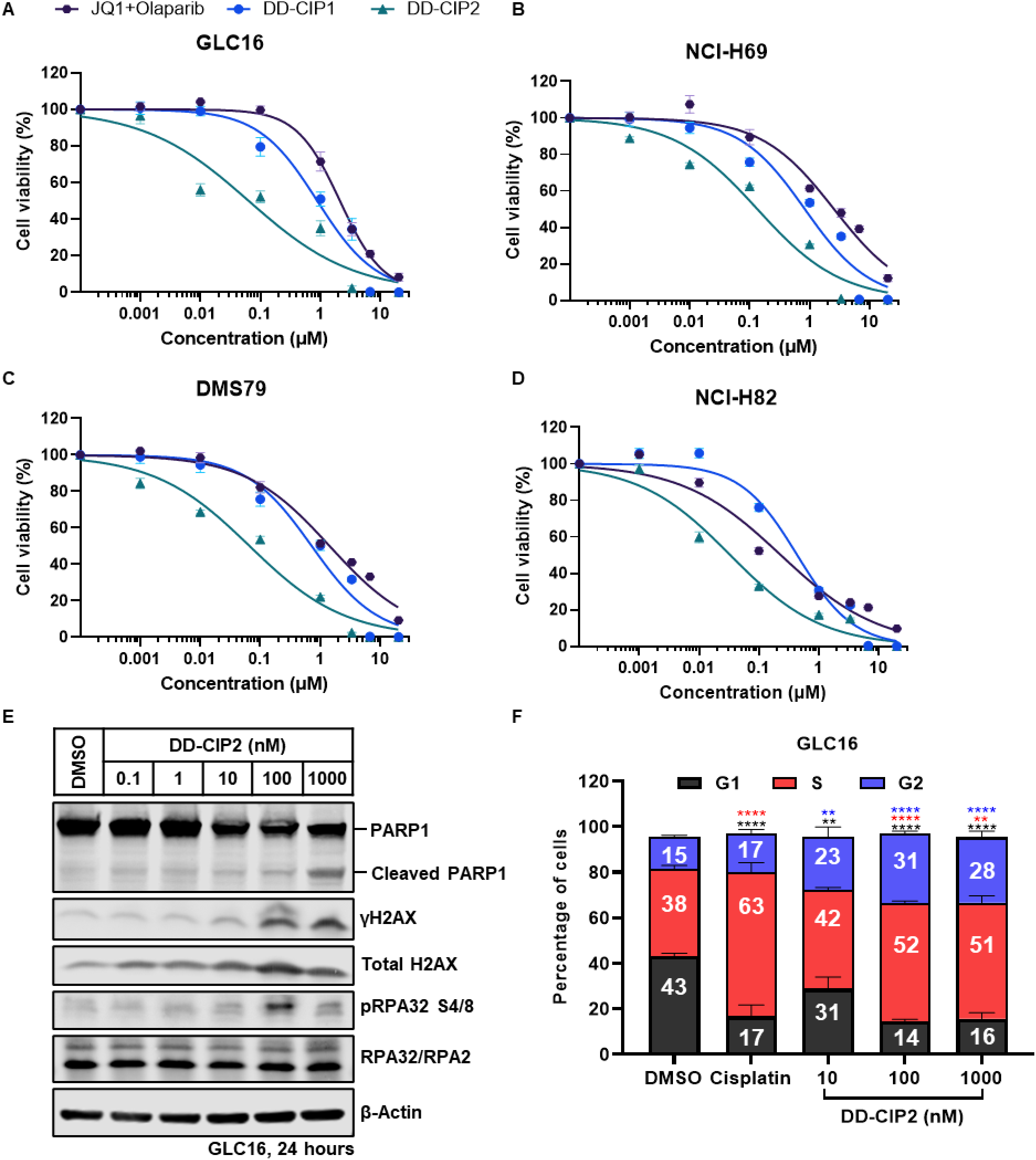
DD-CIP2 effectively induces DNA damage response and anti-tumor efficacy in SCLC model *in vitro*. A-D) Cell-killing potencies of DD-CIP1, DD-CIP2 and co-treatment of JQ1 and Olaparib (1:1) after 72 hours treatment in SCLC cells including A) GLC16; B) NCI-H69; C) DMS79; D) NCI-H82. Data are shown as means ± SEM; n = 3 biological replicates. E) Western blots showing apoptosis (cleaved PARP1) and DNA damage signal (γH2AX, pRPA32 S4/8) markers in GLC16 cells treated with the indicated concentrations of DD-CIP2 for 24 hours. β-actin was used as a loading control. F) Cell cycle analysis of GLC16 cells treated with 10 µM Cisplatin (Used as positive control) or DD-CIP2 at indicated concentrations for 24 hours. Data are shown as means ± SEM; n = 3-7 biological replicates. Statistical significance: **p < 0.01 ***p < 0.001, ****p < 0.0001, one-way ANOVA with Bonferroni’s post-test.

In both GLC16 and NCI-H82 cell lines, DD-CIP2 led to robust induction of DNA damage response, as indicated by elevated levels of γH2AX and cleaved PARP1 in Western blot experiments (**Figure 5E, S7A**). Beyond canonical DNA damage signaling, DD-CIP2 also increased the phosphorylation of RPA32 at Ser4/Ser8, a marker of sustained ATR activation typically associated with DNA end resection inhibition and replication stress checkpoint engagement^52^ (**Figure 5E, S7A**). DD-CIP2 also provoked a dose-dependent accumulation of cells in the G2/M phase, revealed by cell cycle analysis, indicating that DD-CIP2 effectively stalled cell cycle progression 24 hours post treatment (**Figure 5F, S7B-D**). Collectively, these findings demonstrated that DD-CIP2 elicits a coordinated cellular response in SCLC cells involving double-strand break signaling, checkpoint activation, and apoptosis.

We next investigated the anti-tumor efficacy of DD-CIP2 in a human SCLC xenograft model. A 5-day dose-escalation study revealed that DD-CIP2 induced a dose-dependent increase in γH2AX and cleaved PARP1, confirming target engagement in GLC16 xenograft (**Figure 6A,B**). In addition, DD-CIP2 was well tolerated at doses up to 20 mg/kg, with no significant body weight loss (**Figure S8A,B**). Therefore, 20 mg/kg of DD-CIP2 was selected for the efficacy study **(Figure 6C, S8C,D)**. In the efficacy study, vehicle-treated mice exhibited continuous tumor growth, whereas DD-CIP2-treated mice showed strong tumor regression as early as day 3, with durable suppression maintained throughout the experiment period (**Figure 6D,E, S8E,F**). No obvious toxicity was detected, as indicated by steady body weight in the treatment group (**Figure 6F, S8G**). In addition, necropsy revealed no macroscopic abnormalities in major organs. Together, these results demonstrated that DD-CIP2 achieves potent and well-tolerated anti-tumor activity *in vivo* with no apparent systemic toxicity, supporting its potential as a therapeutic strategy for SCLC.

**Figure 6.**
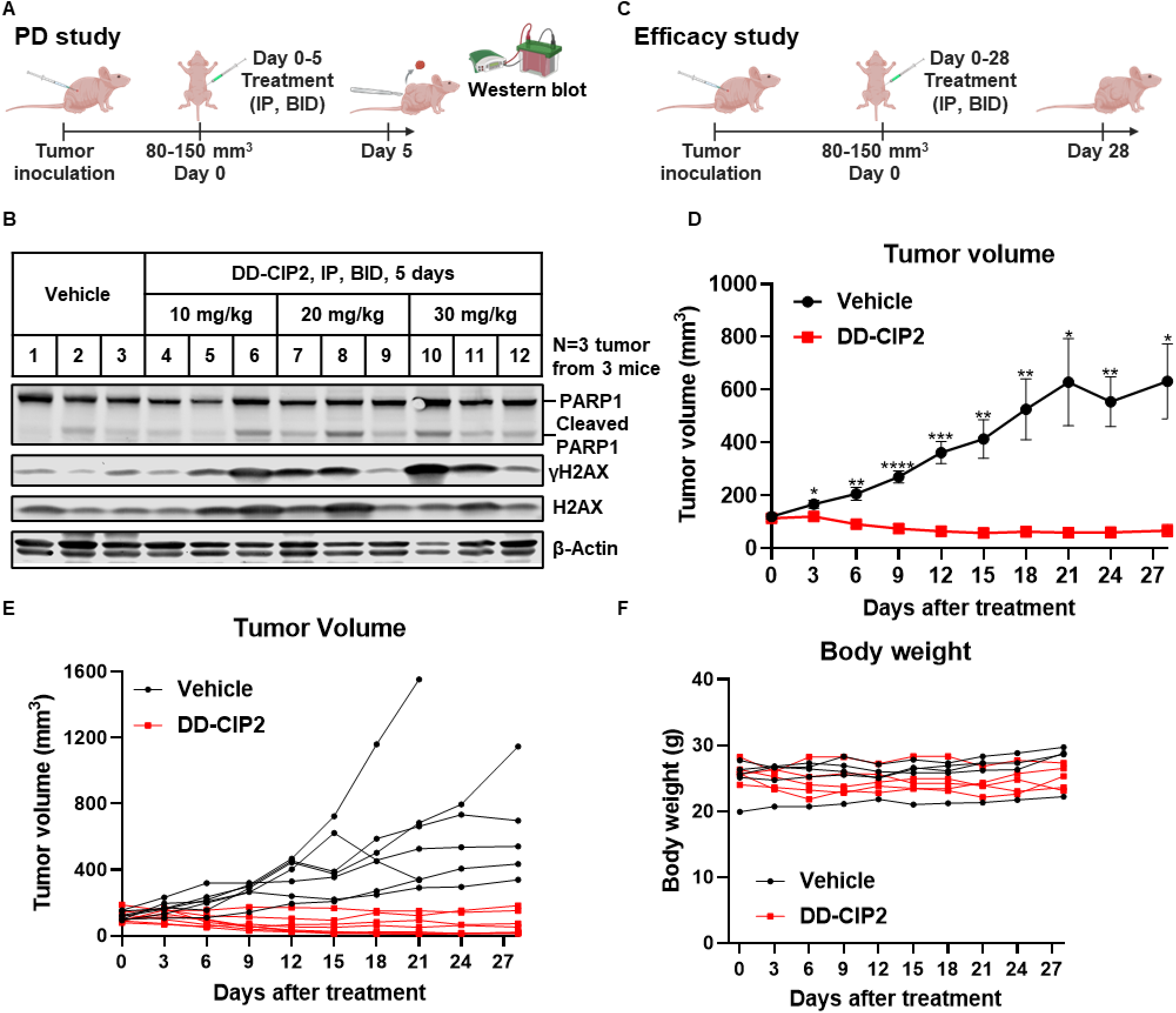
DD-CIP2 effectively induces DNA damage responses and suppresses tumor growth in SCLC model *in vivo*. A) Schematic diagram outlining the dosing strategy of pharmacodynamic (PD) study. B) Western blots showing apoptosis (cleaved PARP1) and DNA damage signal (γH2AX) markers in GLC16 xenograft tumors isolated from the mice after 5 days treatment of vehicle or indicated dosage of DD-CIP2. Each number indicates individual mice. C) Schematic diagram outlining the dosing strategy of efficacy study. D-E) Tumor growth in nude mice injected intraperitoneally (IP) with 20 mg/kg DD-CIP2 twice per day for 28 days. The average (D) tumor volumes are shown as means ± SEM; and the individual tumor growth curve is shown as spider plot (E). Statistical significance: *p < 0.05, **p < 0.01, ***p < 0.001, ****p < 0.0001, with Student’s t-test; n = 7-8 tumor samples per group. (F) Individual spider plots for the body weight.

## Discussion

In this study, we report the discovery and characterization of DNA damage response chemical inducers of proximity (DD-CIPs) that induces BRD4 recruitment to PARP1/2. Unlike conventional occupancy-driven inhibitions, DD-CIPs appear to function through event-driven pharmacology that requires a ternary complex between PARP1/2, DD-CIP1/2 and BRD4, resulting in rapid DDR activation and subsequent apoptosis.

Our medicinal chemistry campaign underscores the importance of both binder selection and linker architecture in optimizing DD-CIP activity. This is a recurrent observation in bivalent small molecule design projects^29^. Specifically, we found that bivalent compounds based on Olaparib with an amide linkage exhibited the most potent anti-proliferative activity. Only specific structural combinations of linkers and ligands produced potency that exceeded co-treatment with the parental inhibitors. The tight dependence on linker length (**Figure 1A**) demonstrates that establishing structure–activity relationships through systematic binder and linker optimization is critical to maximizing proximity-driven pharmacology. Further development of DD-CIPs will entail exploration of alternative ligands for BRD4 and other chromatin regulators^17, 28–30^.

Linker rigidification yielded DD-CIP2, which exhibits potent cytotoxicity across blood cancer and solid tumor cell lines irrespective of BRCA status while showing low cytotoxicity to the human foreskin fibroblast cell line BJ, human lung fibroblast cell line MRC-5, human primary peripheral blood mononuclear cells (PBMC) and the mouse embryonic fibroblast cell line NIH-3T3, yielding a therapeutic index greater than 10 (compared to GLC16 and NCI-H82) and suggesting a low likelihood of toxicity in normal tissues^53–55^. We further demonstrated that DD-CIP2 induces robust DNA damage signaling, disrupts cell cycle progression, and promotes apoptosis in SCLC, a malignancy characterized by profound genomic instability, replication stress, and prevalent loss of RB1 and TP53^56^. These features render SCLC highly dependent on DDR pathways for survival, making it an ideal setting to test DDR-modulating therapeutics^57, 58^. Clinically, SCLC remains as an aggressive disease with limited treatment options. Many cases initially respond to platinum-based chemotherapy, but rapid acquisition of resistance often leads to poor patient outcomes. In this study, we demonstrated that DD-CIP2 induces tumor regression at a well-tolerated dosage in GLC16 xenograft model, underscoring the therapeutic promise of this approach. Establishing proof-of-concept in this context not only validates the mechanistic potential of DD-CIPs to exploit DDR vulnerabilities but also highlights their translational relevance in a cancer type with a high unmet therapeutic need. More broadly, DD-CIPs represent a mechanistically distinct pharmacology that enhances and extends the efficacy of PARPi, even in HR-proficient cancers.

While we were preparing our manuscript, da Camara *et al.* reported PCIPs, the bivalent compounds composed of JQ1, Rucaparib and variable linkers^59^. They identified PARP2 as the relevant isoform mediating the biological response of PCIP1, a compound that shares structural similarity with TWQ-03-073. Similarly, DD-CIP1 and DD-CIP2 induced binding between BRD4 and both PARP1 and PARP2. While a PARP1/2 dual inhibitor (Veliparib and Olaparib) effectively reversed the cellular cytotoxicity of DD-CIP1 and DD-CIP2, the specific contributions of each PARP isoform remain to be fully defined. Future genetic perturbation and the use of isoform-selective ligands will be valuable to clarify which PARP isoform(s) are essential for DD-CIP activity and how their engagement influences downstream DNA damage response.

Given these findings, we hypothesize that compound-induced PARP1/2-BRD4 binding could establish a positive feedback loop that amplifies DNA damage stress. Like PARP inhibitors, DD-CIPs likely exploit elevated DNA damage in cancer cells and, even more than monovalent inhibitors, further intensify genotoxic stress, resulting in potent activity in tumor cells while exerting limited effects on normal cells^3, 60–63^. The strong tumor-selective activity of DD-CIPs may reflect aberrant mislocalization of BRD4 on chromatin^64^ or mislocalization of PARP1. Both molecular perturbations would compromise the timely recruitment of DNA repair factors and attenuate the resolution of otherwise innocuous DNA lesions^65^. Another intriguing possibility is that BRD4 may drive aberrant transcription at DNA damage sites, where transcriptional repression is normally required to facilitate error-free repair^66, 67^. Further mechanistic dissection is required to test this model and to inform the rational design of next-generation DD-CIP compounds.

## Methods

### General cell biology methods

Roswell Park Memorial Institute (RPMI) 1640 medium and Dulbecco’s modified Eagle’s medium (DMEM),, Heat-inactivated Fetal bovine serum (FBS), penicillin-streptomycin (10,000 units/mL sodium penicillin G and 10,000 μg/mL streptomycin), trypsin-EDTA solution (1×), and phosphate-buffered saline (PBS; 1×) were purchased from Gibco Invitrogen Corp (Grand Island, NY, USA). Eagle’s Minimum Essential Medium (EMEM) and DMSO were purchased from ATCC (Manassas, Virginia, USA). dBET6, Olaparib, Veliparib and Talazoparib were purchased from MedChemExpress (Monmouth Junction, NJ, USA). All other chemicals were purchased from Sigma-Aldrich (St. Louis, MO, USA), unless indicated otherwise.

### Cell lines

HEK293T, Jurkat, MOLT-4, SU-DHL-5, MDA-MB-231, NCI-H2452, HepG2, A549, NCI-H3122, NCI-H460, GLC16, NCI-H69, NIH-3T3, DMS79, BJ (human fibroblasts, CRL-2522), MRC-5, human PBMC (PCS-800-011, Lot: 80413251) cells were purchased and verified from the ATCC. NCI-H82 cells were a gift from the laboratory of John T. Poirier (NYULH) and originally from the ATCC. KARPAS422 cells were obtained from Sigma (06101702). Daudi cells were a gift from the laboratory of R. Levy (Stanford University) and originally from the ATCC. Raji cells were a gift from the laboratory of J. Cochran (Stanford University) and originally from the ATCC. PANC-1 cells were a gift from the laboratory of S. Corsello (Stanford University) and originally from the ATCC. MDA-MB-436, MDA-MB-468, TYK-nu cells were a gift from the laboratory of P. Sorger (Harvard Medical School). HEK293T, MDA-MB-231, A549, MDA-MB-436, MDA-MB-468 and NIH-3T3 cells were cultured in DMEM medium, with 10% heat-inactivated FBS and 1% penicillin-streptomycin. Jurkat, MOLT-4, SU-DHL-5, Daudi, Raji, TYK-nu, PANC-1, NCI-H2452, NCI-H3122, NCI-H460, GLC16, NCI-H82, NCI-H69, DMS79 and human PBMC cells were cultured in RPMI medium, with 10% FBS and 1% penicillin-streptomycin. HepG2, BJ (human fibroblasts) and MRC-5 cells were cultured in EMEM medium, with 10% FBS, 1% penicillin-streptomycin. All cell lines were grown in humidified tissue culture incubators at 37 ℃ with 5% CO_2_ and were mycoplasma-negative based on monthly testing using the MycoAlert mycoplasma detection kit (Lonza, Basel, Switzerland). For all experiments, cells had undergone fewer than 18 passages.

### PARP1 NanoBRET assay for target engagement

The reagents for PARP1 NanoBRET assay (CS3489A26, CS3489A07) were purchased from Promega through Academic Access Program (AAP). And the assay was performed based on the manufacture.

HEK293T cells were trypsinized and brought to a concentration of 2×10e5 cells/mL in Opti-MEM with 1% FBS. 21 mL of cells in above density was transfected with transfection mixture containing 1.15 μg PARP1-Nanoluc fusion vector (Promega # CS3489A07), 10 μg Transfection Carrier DNA (Promega # E4881), 32 μL FuGENE HD transfection reagent (Promega #E2311) and 1 mL Opti-MEM. The cells were directly plated into 384 well plates384-well, white, tissue culture treated, opaque bottom plates, with 44 μL/well volume. The transfected cells are allowed to incubate for 24 hours at 37 °C. Following the 24 hours incubation, the NanoBRET® Tracer PARP-01 (Promega # CS3489A26) solution was then prepared by diluting the stock PARP-01 tracer with DMSO to make a 100X stock. The 100X K-8 stock was then further diluted using Promega dilution buffer to prepare a 20X stock. The 20X stock was further diluted using Opti-MEM to make an 8.33X stock. 6 μL of the 8.33X PARP-01 stock was dispensed into each well for a final concentration of 12.5 nM PARP-01 tracer in cells. 6 μL of Opti-MEM were added to designated wells in place of the PARP-01 tracer solution to act as no acceptor controls for data analysis. The plate was then treated with compounds using a Tecan D300e digital dispenser and allowed to incubate for indicated time at 37 °C. After incubation, a 3X NanoBRET NanoGlo substrate with extracellular NanoLuc inhibitor solution was made, and 20 μL was dispensed into each well. The plate was mixed for 30 s at 350 RPM on an orbital shaker. Data was then obtained using a PheraStar plate reader measuring luminescence with 450-BP and 610-LP filters.

### HiBiT-BRD4 assay for BRD4 target engagement

The generation of HiBiT-BRD4 Jurkat cells was reported previously^42^. Endogenous BRD4 protein levels were evaluated using the Nano-Glo HiBiT Lytic Detection System (Promega). In brief, 1.8 × 10^4^ HiBiT-BRD4 Jurkat cells were seeded into 384-well plates and incubated with the indicated concentrations of compounds for indicated time. Then the plates were subjected to the treatment of 0.5 μM of dBET6 for 45 minutes. After incubation, the plates were subjected to Nano-Glo HiBiT Lytic Detection System as described in manufacturer’s manual. The HiBiT-BRD4 assays were conducted in biological triplicates.

### NanoBRET assay for PARP1-BRD4 ternary complex

HEK293T cells were plated at a density of 2×10e5 cells/mL in 2mL of DMEM/well in a tissue culture treated, 6 well plate and were allowed to incubate overnight at 37 °C. The next day, each well of cells was transfected with 0.01 μg NanoLuc-BRD4 and 1 μg HaloTag-PARP1 using FuGENE HD transfection reagent (Promega #E2311) for a ratio of 1:100 NanoLuc to HaloTag plasmids. The transfected cells were allowed to incubate overnight at 37℃. The following day, the transfected cells were trypsinized and brought to a concentration of 2×10e5 cells/mL in Fluorobrite DMEM containing 10% FBS. 40uL of cells per well were plated in a 384 well plate and treated with 100 nM HaloTag618 Ligand. Designated wells were not treated with the HaloTag618 Ligand and normalized with DMSO to be used as no acceptor controls for data analysis. The cells were allowed to incubate overnight at 37℃. The next day, the cells were treated with compounds for testing using a Tecan D300e and incubated at 37℃ for 2 hours. After 2 hours, 10 μL of 5x NanoBRET NanoGlo Substrate was added to each well and mixed on an orbital shaker at 350 RPM for 30 sec. Data was then obtained using a PheraStar plate reader measuring luminescence with 450-BP and 610-LP filters.

For all nanoBRET assays, a dose-response curve with 3 technical replicates for each compound carried out, and corrected BRET ratios were calculated according to manufacturer assay protocol (Promega #TM439). Data was fit using a standard four parameter (variable slope) function using GraphPad Prism 10.

### Cell viability analysis

Cells were seeded in 96 or 384-well plates in media and treated with indicated compounds at various concentrations for 72h. Cell viability was measured using the CellTiter-Glo® 2.0 Cell Viability Assay (G9242, Promega, USA) except for BJ (human fibroblasts) cells. Data was fit using a standard four parameter (variable slope) function using GraphPad Prism 10.

Jurkat, KARPAS422, Daudi, Raji: 384 well, 500 cells/well

MOLT-4, SU-DHL-5, MRC-5: 384 well, 1000 cells/well

MDA-MB-436, MDA-MB-468, TYK-nu, MDA-MB-231, NCI-H2452, HepG2, A549, NCI-H3122, NCI-H460, PANC-1: 384 well, 500 cells/well

GLC16, NCI-H82, NCI-H69, DMS79 and NIH-3T3: 96 well, 2500 cells/well

PBMC: 384 well, 5500 cells/well

#### Cell viability in BJ (human fibroblasts) cells

5000 cells were seeded in 100 µl of media per well of a 96-well plate and treated with drug for the indicated times and doses. A resazurin-based indicator of cell health (PrestoBlue; P50200, Thermo Fisher) was added for 1.5 h at 37°C, at which point the fluorescence ratio at 560/590 nm was recorded (Tecan Spark). The background fluorescence was subtracted, and the signal was normalized to DMSO-treated cells.

### Western blot analysis

The cells were washed once with DPBS before lysing in RIPA lysis buffer (50 mM Tris, 150 mM NaCl, 0.5% deoxycholate, 0.1% SDS, 1.0% Triton X-100, pH 7.4) (BP-116TX, Boston Bioproducts, Inc., Milford, MA, USA) supplemented with a protease inhibitor cocktail (11836153001, Roche) for 15 min on ice. The lysates were centrifuged at 13,000g (4 °C, 20 minutes), and the supernatant was collected. Protein concentrations of the cell lysates were quantified using the BCA method and a BCA Protein Assay Kit (Thermo Fisher Scientific, Waltham, MA, USA). Equal amounts of protein were mixed with 6x Laemelli sample buffer (J61337.AC, Thermo Fisher Scientific) containing 150 mM DTT and then subjected to 4-20% sodium dodecyl sulfate polyacrylamide gel electrophoresis (SDS-PAGE) and transferred to nitrocellulose membranes (cat. no. 1620112; Bio-Rad Laboratories, Hercules, CA, USA). The membranes were blocked using Intercept® (TBS) Blocking Buffer (LI-COR Biosciences, Lincoln, NE, USA) and subsequently probed with appropriate primary antibodies [anti-phospho-ATM S1981 (cat. no. 13050; Cell Signaling Technology, Danvers, MA, USA), anti-phospho-ATR S428 (cat. no. 2853; Cell Signaling Technology, Danvers, MA, USA) anti-phospho-CHK1 S345 (cat. no. 2348; Cell Signaling Technology, Danvers, MA, USA) anti-phospho-CHK2 T68 (cat. no. 2197; Cell Signaling Technology, Danvers, MA, USA), anti-BRD4 (cat. no. ab128874; Abcam, Cambridge, UK), anti-α-Tubulin (cat. no. 3873; Cell Signaling Technology, Danvers, MA, USA), anti-FLAG M2 (cat. no. F1804; Sigma-Aldrich), anti-RPA32/RPA2 (2208, Cell Signaling Technology, Danvers, MA, USA), anti-phospho-RPA32 S4/S8 (A700-009-T, Bethyl Laboratories, USA) anti-phospho-histone H2A.X (9718, Cell Signaling Technology, Danvers, MA, USA), anti-histone H2A.X (2595S, Cell Signaling Technology, Danvers, MA, USA), anti-PARP1 (9542, Cell Signaling Technology, Danvers, MA, USA), anti-beta actin(A2228, Sigma-Aldrich, USA)] at 4 °C overnight. The membranes were washed with TBST (20 mM Tris, pH 7.5, 150 mM NaCl, 0.1% Tween-20) (IBB-180X, Boston Bioproducts, Inc., Milford, MA, USA) for 4*5 min at room temperature and incubated with IRDye 800-labeled goat anti-rabbit IgG (LI-COR Biosciences, cat. no. 926-32211) or IRDye 680RD goat anti-Mouse IgG (LI-COR Biosciences, cat. no. 926-68070) secondary antibodies at room temperature for 1 hour. After washing the membranes with TBST for 4*5 min at room temperature, the membranes were detected on Li-COR Odyssey CLx system.

For the immunoprecipitation samples, HEK293T cells were transfected with pcDNA5-Flag-BRD4-WT (Addgene #90331) or pCMV-PARP1-3xFlag-WT (Addgene #111575) using Lipofectamine 2000 following manufacture instructions. After 36h, the cells were treated with indicated compounds for certain time, and the cell lysates were prepared in NP-40 IP lysis buffer (50 mM Tris, 150 mM NaCl, 5 mM EDTA, 1% Nonidet P-40, pH 7.4) (BP-119, Boston Bioproducts, Inc., Milford, MA, USA) supplemented with a protease inhibitor cocktail and then centrifuged at 20,000g for 15 min at 4 °C. 10% of lysate was saved for the “Input” and the rest of lysates were incubated with 10 μL Anti-FLAG® M2 magnetic beads (Millipore-sigma) overnight at 4 °C with gentle rotation. The beads were washed once with NP-40 IP lysis buffer and twice with 0.02% Tween-20 in PBS to remove nonspecific proteins. The enriched protein was eluted with 30 μL of 50 mM glycine (pH 2.8) for 5 min at room temperature and brought to neutral with 2 μL of 1 M Tris-base, “output”. Both input and output protein were mixed with 6x Laemelli sample buffer (Alfa Aesar) containing 150 mM DTT, then analyzed by SDS-PAGE and Western Blot.

### Immunoprecipitation-Mass Spectrometry (IP-MS)

Immunoprecipitation was performed as described above in three biologically independent replicates, each replicate contains 1 mg total protein and 50 µL Anti-FLAG® M2 magnetic beads. The washing buffer is modified as: 1) wash 4 x 1 mL with IP lysis buffer; 2) wash 3 x 1 mL with 25 mM Tris, 150 mM NaCl; 3) wash 3 x 1 mL with MS grade water. Enriched proteins were eluted three times with 150 µL of 3% v/v ammonium hydroxide (pH 11-12) for 30 min with gentle rotation at cold room. The collected eluate was then dried using a centrifugal vacuum concentrator (Thermo SPD120-115). The samples were reconstituted in 50 µL of reduction-alkylation buffer containing 8 M urea, 10 mM TCEP, 40 mM chloroacetamide, 4.4 mM CaCl_2_, and 100 mM HEPES, pH 8.0 and then incubated for at least 30 min while shaking. The samples were then diluted with 160 µL of 20 mM HEPES, pH 8.0, and then digested overnight at 37 °C with 300 ng of MS-grade Trypsin/Lys-C Protease Mix (final concentration of 1.33 ng/µL). The samples were then acidified by adding trifluoroacetic acid (TFA) until the sample reached pH ≤ 3, as confirmed by pH paper.

*SPE Desalting.* Samples were then desalted using SOLAµ SPE HRP Peptide Desalting columns (Thermo 60209-001) with a Positive Pressure Manifold and all MS-grade reagents. The desalting cartridges were activated with 200 µL of acetonitrile and then equilibrated with two washes of 200 µL Wash Buffer (2% ACN + 0.2% TFA in water). After loading samples into the cartridges, the samples were washed three times with 200 µL Wash Buffer and then eluted using 100 µL 50% acetonitrile with 0.1% formic acid (FA) in water. The desalted peptides were then immediately dried on a centrifugal vacuum concentrator (Thermo SPD120-115). The samples were then reconstituted in 0.1% FA in water and then analyzed by LC-MS/MS (see below).

### LC-MS/MS diaPASEF Acquisition & Statistical Analysis

#### diaPASEF LC-MS/MS Analysis

The reconstituted desalted peptides were then analyzed using a nanoElute 2 UHPLC (Bruker Daltonics, Bremen, Germany) coupled to a timsTOF HT (Bruker Daltonics, Bremen, Germany) via a CaptiveSpray nano-electrospray source. The peptides were separated in the UHPLC using an Aurora Ultimate nanoflow UHPLC column with CSI fitting (25 cm x 75 µm ID, 1.7 µm C18; IonOptics AUR3-25075C18-CSI) over a 70 min gradient at a flow rate of 400 nL/min with column temperature maintained at 50 °C using Mobile Phase A (MPA; 3% acetonitrile + 0.1% FA in water) and Mobile Phase B (MPB; 0.1% FA in ACN). The 70 min gradient started at 6% MPB while increasing to 17% MPB at 40 min, 25% MPB at 55 min, 34% MPB at 64 min, 85% MPB at 65 min, and finishing at 92% MPB at 70 min.

The TIMS elution voltages were calibrated linearly with three points (Agilent ESI-L Tuning Mix Ions; 622, 922, 1,222 m/z) to determine the reduced ion mobility coefficients (1/K_0_). diaPASEF was performed using the MS settings 100 m/z for Scan Begin and 1700 m/z for Scan End in positive mode, the TIMS settings 0.70 V⋅s/cm^2^ for 1/K_0_ start, 1.30 V⋅s/cm^2^ for 1/K_0_ end, ramp time of 120.0 ms, 100% duty cycle, ramp rate of 7.93 Hz, and the capillary voltage set to 1600 V. diaPASEF windows from mass range 226.8 Da to 1226.8 Da and mobility range 0.70 1/K_0_ to 1.30 1/K_0_ were designed to provide 25 Da windows covering doubly and triply charged peptides as confirmed by DDA-PASEF scans, whereas singly charged peptides were excluded from the acquisition due to their position in the m/z-ion mobility plane.

#### Raw data processing

The raw diaPASEF files were processed using library-free analysis in FragPipe (22.0 or 23.1). DIA spectrum deconvolution was performed using diaTracer (1.1.5 or 1.3.5) with the following default settings: (i) “Delta Apex IM” to 0.01, (ii) “Delta Apex RT” to 3, (iii) “RF max” to 500, (iv) “Corr threshold” to 0.3, and (v) mass defect filter enabled with offset set to 0.1^68, 69^. The reviewed Homo sapiens protein sequence database was obtained from UniProt (07/13/2024; 20,468 entities) with decoys and common contaminants. In the MS Fragger (4.1 or 4.3) database search, the following settings were used: (i) initial precursor and fragment mass tolerances of 10 ppm and 20 ppm, respectively, (ii) enabled spectrum deisotoping, mass calibration, and parameter optimization, and (iii) isotope error set to “0/1/2”. For protein digestion, “stricttrypsin” for fully tryptic peptides was enabled with up to 1 missed cleavage, peptide length from 7 to 50, and peptide mass range from 500 to 5,000 Da. For modifications, methionine oxidation (2 max occurrences) and N-terminal acetylation (1 max occurrence) were set as variable modifications (maximum up to 3), while cysteine carbamidomethylation was set as a fixed modification. For validation, MSBooster (DIA-NN model) and Percolator were used for RT and MS/MS spectra prediction and PSM rescoring, while ProteinProphet (--maxppmdiff 2000000) was used for protein inference with FDR filtering (--picked --prot 0.01). The spectral library was then generated by EasyPQP 0.1.49 using default settings. Peptides were then quantified using DIA-NN (1.9.1 or 2.2.0 Academia) with 0.1% FDR and QuantUMS high precision settings. The DIA-NN parquet report containing peptide quantification and scoring was then analyzed in R.

#### Statistical Analysis

DIA-NN quantified peptides were then further filtered by (i) removing common contaminants and decoy sequences, (ii) 1% FDR filtering at the global.q.value (precursor Q value across all samples) and pg.q.value (protein group Q value in single injection), (iii) removing peptides which were not detected in at least two replicates in a single condition, and (iv) removing singly charged peptides and non-proteotypic (unique) peptides. Protein intensities were then re-calculated using the MaxLFQ method provided in the DIA-NN R package^70^. Differential statistics was then performed using the DEqMS R package^71^, which performs a LIMMA-moderated t-test with an adjustment for number of detected peptides per protein, to determine the p-value, fold change, and Benjamini-Hochberg adjusted p-value.

### Cell cycle analysis

Cells were seeded in 6-well plates at a density of 1 × 10^6^ cells per well and treated with DD-CIP2 at the indicated concentrations for 24 hours. Following treatment, cells were harvested by centrifugation, washed with cold PBS, and fixed in 1 ml of 75% ethanol in PBS overnight at – 20 °C. Fixed cells were then collected by centrifugation, washed twice with cold PBS, and resuspended in FxCycle™ PI/RNase Staining Solution (F10797, Invitrogen, USA). Samples were incubated for 15 min at room temperature in the dark before analysis by flow cytometry. Cell populations were gated based on PI staining, and the distribution across G1, S, and G2/M phases was quantified using FlowJo software.

### Mouse pharmacokinetic (PK) study

The PK study was performed in the Drug Metabolism and Pharmacokinetics (DMPK) Core facility at Scripps Florida (https://www.scripps.edu/science-and-medicine/coresand-services/dmpk-core/index.html). The mice are housed in individually ventilated cages (IVC) in JAG 75 cages with micro-isolator lids. HEPA filtered air is supplied into each cage at a rate of 60 air exchanges per hour for the mice. The dark/light cycle is set for 8:00pm on-8:00pm off. The temperature is set for 72 OF and is maintained +/-2 OF. The humidity is low/Hi of 30-70%. There is a computerized system in place to control and or monitor the temperatures within the Animal Holding Room. Each animal room is equipped with a thermos-hygrometer that is monitored and recorded daily on room log. 3 mg/kg (Intravenous dosing, IV) or 10 mg/kg (Intraperitoneal dosing, IP) of DD-CIP2 were injected into C57BL/6 male mice (n=3) using a 25-29 gauge needle to deliver 10 μL/g body weight of a formulation of 0.5 mg/mL DD-CIP2 in 5% DMSO and 95% Saline. The formulation was checked to be a clear solution and after administration, the animal was put back in its cage. Plasma levels were measured at 0, 5, 15, 30, 60, 120, 240, 360, and 480 mins after drug administration. Samples were processed for analysis by precipitation using acetonitrile and analyzed with LC/MS/MS. PK parameters were calculated using the noncompartmental analysis tool of WinNonlin Enterprise software (version 6.3). All procedures were approved by the Scripps Florida Institutional Animal Care and Use Committee (IACUC), and the Scripps Vivarium is fully accredited by the Association for Assessment and Accreditation of Laboratory Animal Care International.

### Animal studies

All animal procedures were reviewed and approved by the Institutional Animal Care and Use Committee (IACUC) at the New York University Langone Health (NYULH). NU-Foxn1nu (Nude, Cat#: 088) mice were used for SCLC xenograft studies. Animals were housed and maintained in accordance with respective NYULH and CAS for the care, welfare, and treatment of laboratory animals. All experimental procedures complied with, or exceeded, the standards of the Association for the Assessment and Accreditation of Laboratory Animal Care, International (AAALAC), the United States Department of Health and Human Services, and applicable local and federal animal welfare laws. At 8 weeks of age, mice were subcutaneously inoculated with 2 × 10^6^ GLC16 cells into both flanks. Tumor dimensions (length, width and height) were measured using calipers, and tumor volumes were calculated with the formula: Volume = (π/6) × Length × Width × Height^72–74^. Once tumors reached 80-150 mm^3^, mice were randomized into treatment groups and administrated either vehicle control (5% DMSO, 5% Tween 80, 40% PEG300, 50% Saline), or DD-CIP2 at 20 mg/kg (I.P., BID). Tumor size and body weight were recorded every 3 days.

### Chemical synthesis

Synthesis details are provided in the Supplementary Methods (Chemical synthesis).

## Data availability

Data generated in this study are provided in the manuscript, supplementary information, and source data files. Additional data supporting the findings of this study are available from the corresponding author upon reasonable request.

## Acknowledgements

This work was supported by the National Institutes of Health (NIH) grants RO1 5R01CA282437 (to K.K.W.), CA276167, CA163915, and MH126720-01 (to G.R.C.), NIH High End Instrumentation grant (1S10OD028697-01) (to N.S.G.), departmental funds from Stanford Chemical and Systems Biology and Stanford Cancer Institute (to N.S.G.). Funding for pharmacokinetic studies was provided by NIH grant number 1 S10OD030332-01. G.R.C. received support for this work from the Spark Foundation (SPO #349451) at Stanford and the generous support of the Victor Family Foundation. PARP1 NanoBRET assay was supported by Promega Corporation through Academic Access Program (AAP). Y.L.W was supported by the Lucile Packard Foundation for Children Health, Coxe-Otus Fellowship Research Foundation.

