## Supplementary Figures for "Design and Development of DNA Damage Chemical Inducers of Proximity (DD-CIP) for Targeted Cancer Therapy"

**A**

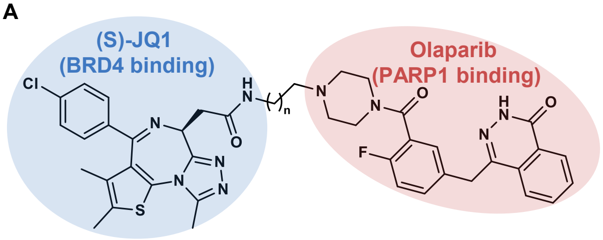


| Compound ID (TWQ-03-XXX) | n | 72 hours $IC_{50}$ (nM) | |
|---|---|---|---|
| | | MOLT-4 | SU-DHL-5 |
| 086 | 8 | 369 | 108 |

**B**

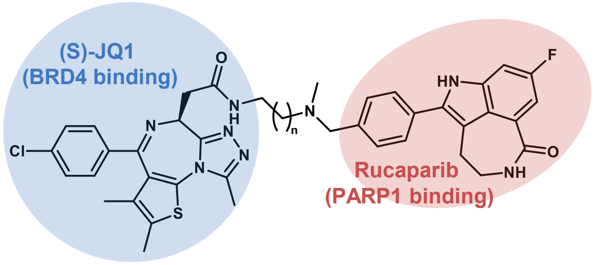


| Compound ID (TWQ-03-XXX) | n | 72 hours $IC_{50}$ (nM) | |
|---|---|---|---|
| | | MOLT-4 | SU-DHL-5 |
| JQ1+Rucaparib | | 208 | 79 |
| 073 | 6 | 113 | 68 |
| 089 | 7 | 141 | 101 |
| 097 | 9 | 515 | 265 |
| 105 | 10 | 548 | 457 |
| 101 | 11 | 928 | 529 |

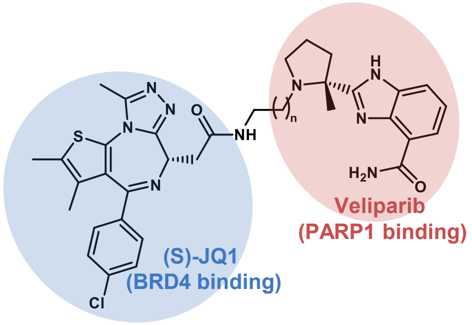


| Compound ID (TWQ-03-XXX) | n | 72 hours $IC_{50}$ (nM) | |
|---|---|---|---|
| | | MOLT-4 | SU-DHL-5 |
| JQ1+Veliparib | | 295 | 146 |
| 072 | 6 | 218 | 125 |
| 087 | 7 | 455 | 218 |
| 095 | 9 | 1339 | 432 |
| 103 | 10 | 920 | 455 |
| 099 | 11 | 4903 | 2586 |

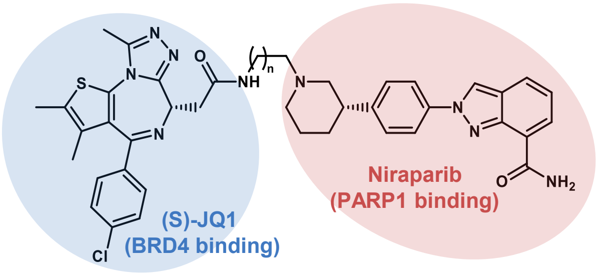


| Compound ID (TWQ-03-XXX) | n | 72 hours $IC_{50}$ (nM) | |
|---|---|---|---|
| | | MOLT-4 | SU-DHL-5 |
| JQ1+Niraparib | | 224 | 118 |
| 071 | 6 | 124 | 104 |
| 088 | 7 | 202 | 116 |
| 096 | 9 | 509 | 325 |
| 104 | 10 | 533 | 461 |
| 100 | 11 | 801 | 538 |

**Figure S1.** Development of DD-CIPs. A) Chemical structure of TWQ-03-086 and cell-killing potencies of TWQ-03-086 at 72 hours in SU-DHL-5 and MOLT-4 cells. Data are shown as means; n = 3 biological replicates. B) Chemical structure of DD-CIPs with different PARP1 ligand and cell-killing potencies of DD-CIP compounds compared to the co-treatment of indicated parental inhibitors at 72 hours in SU-DHL-5 and MOLT-4 cells. Data are shown as means; n = 4 biological replicates.

A

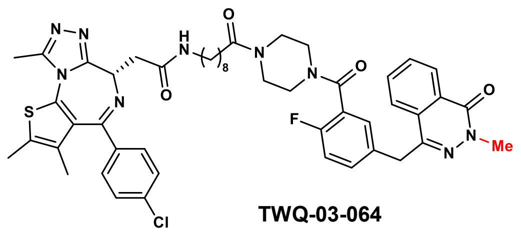


B

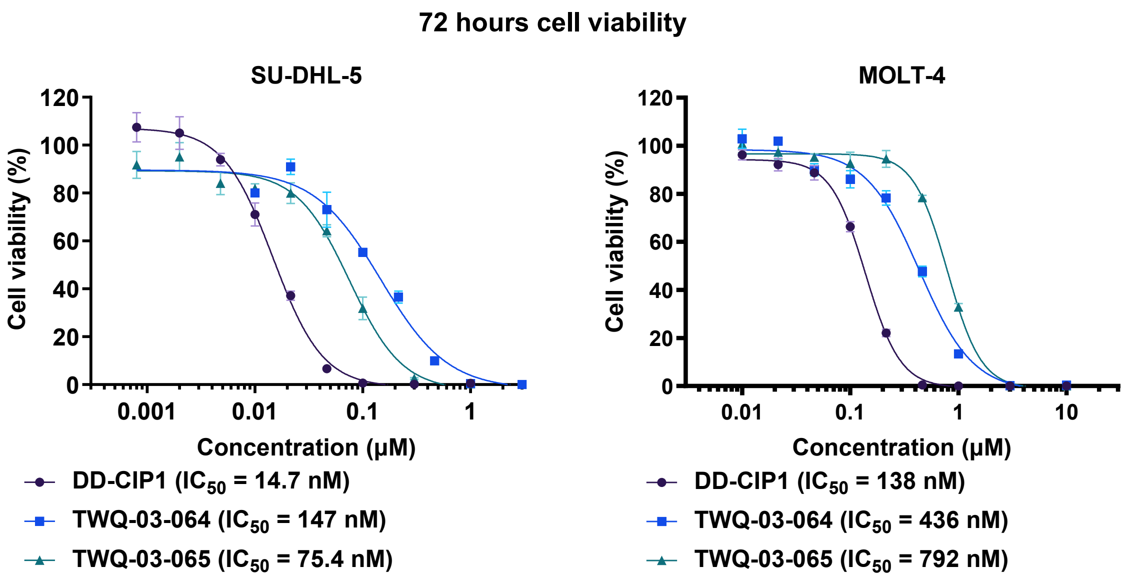


**Figure S2.** Negative compound shows weaker cytotoxicity compared to active compound. A) Chemical structure of TWQ-03-064 and TWQ-03-065. The crucial modification is highlighted with red color. B) Cell-killing potencies of DD-CIP1, TWQ-03-064 and TWQ-03-065 after 72 hours treatment in SU-DHL-5 and MOLT-4 cells. Data are shown as mean ± SEM.; n = 3 biological replicates.

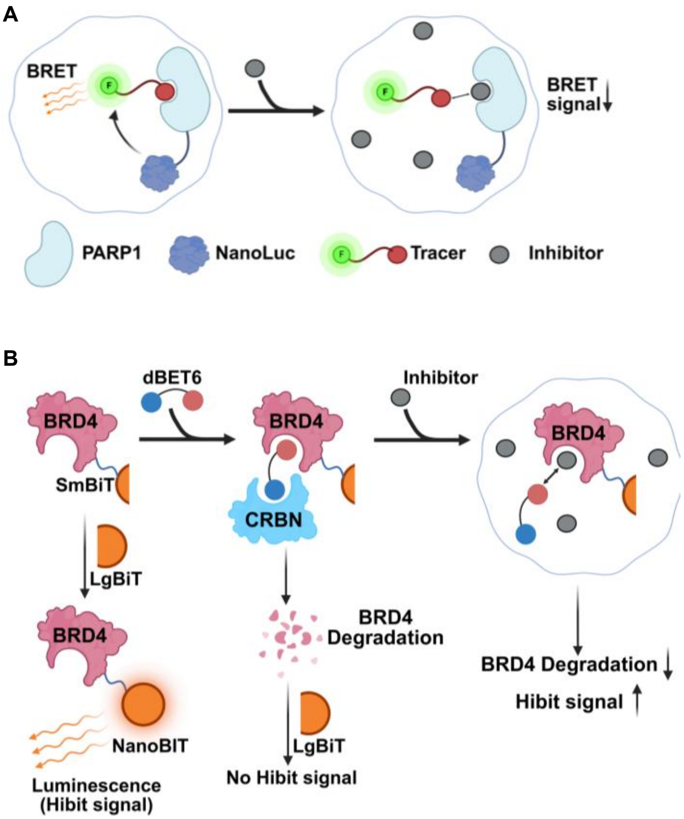


**Figure S3.** Scheme of target engagement assay on: A) PARP1; B) BRD4.

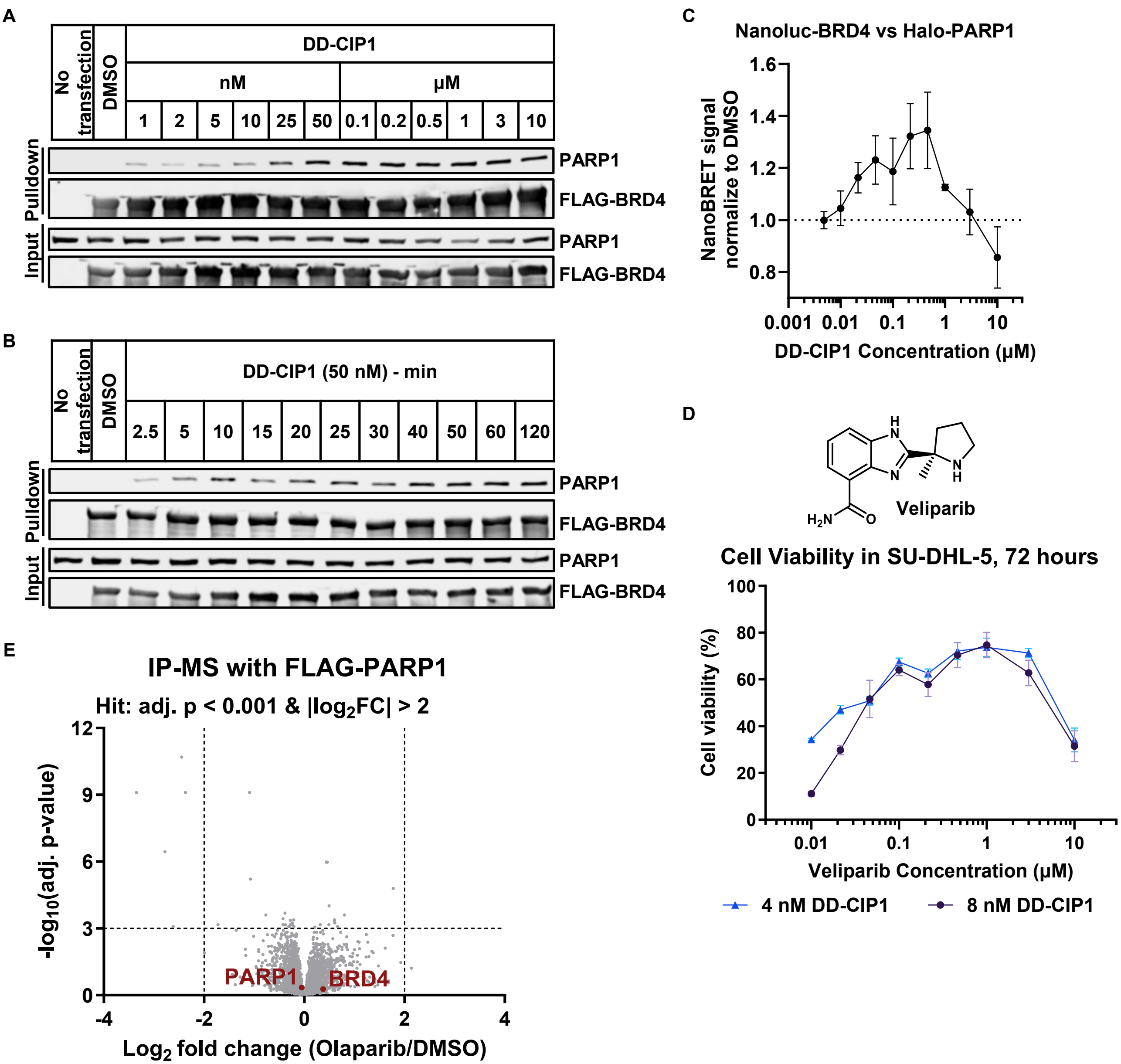


**Figure S4.** DD-CIP1 induces neo-interaction between PARP1 and BRD4. A) Co-IP of FLAG-BRD4 and PARP1 in HEK293T cells after 1 hour treatment of indicated concentrations of DD-CIP1. B) Co-IP of FLAG-BRD4 and PARP1 in HEK293T cells treated with 50 nM DD-CIP1 for indicated time. C) Ternary complex formation between Nanoluc-BRD4 and HaloTag-PARP1 in HEK293T cells after addition of DD-CIP1; means ± SEM, n=3 technical replicates. D) Measurement of SU-DHL-5 cell viability after competitive titration of Veliparib to constant 4 or 8 nM DD-CIP1; cells were treated simultaneously with DD-CIP1 and Veliparib for 72 hours. Data are shown as means ± SEM; n = 3 biological replicates. E) FLAG IP-MS in HEK293T cells overexpressing FLAG-PARP1 treated with 100 nM Olaparib for 1 hour; plotted with cut-offs of $|\log_2(\text{fold change})| \geq 1.25$ and adj. $P \leq 0.001$; 3 biological replicates. All P values were adjusted using Benjamini-Hochberg method from LIMMA-moderated t-test.

| | Mouse microsomal stability | |
|---|---|---|
| **Compound** | **$T_{1/2}$ -min** | **$Cl_{int}$ - µl/min/mg** |
| **sunitinib** | **17.0** | **41** |
| **DD-CIP1** | **2.3** | **307** |

**Figure S5.** Mouse microsome stability of DD-CIP1. Sunitinib is used as reference.

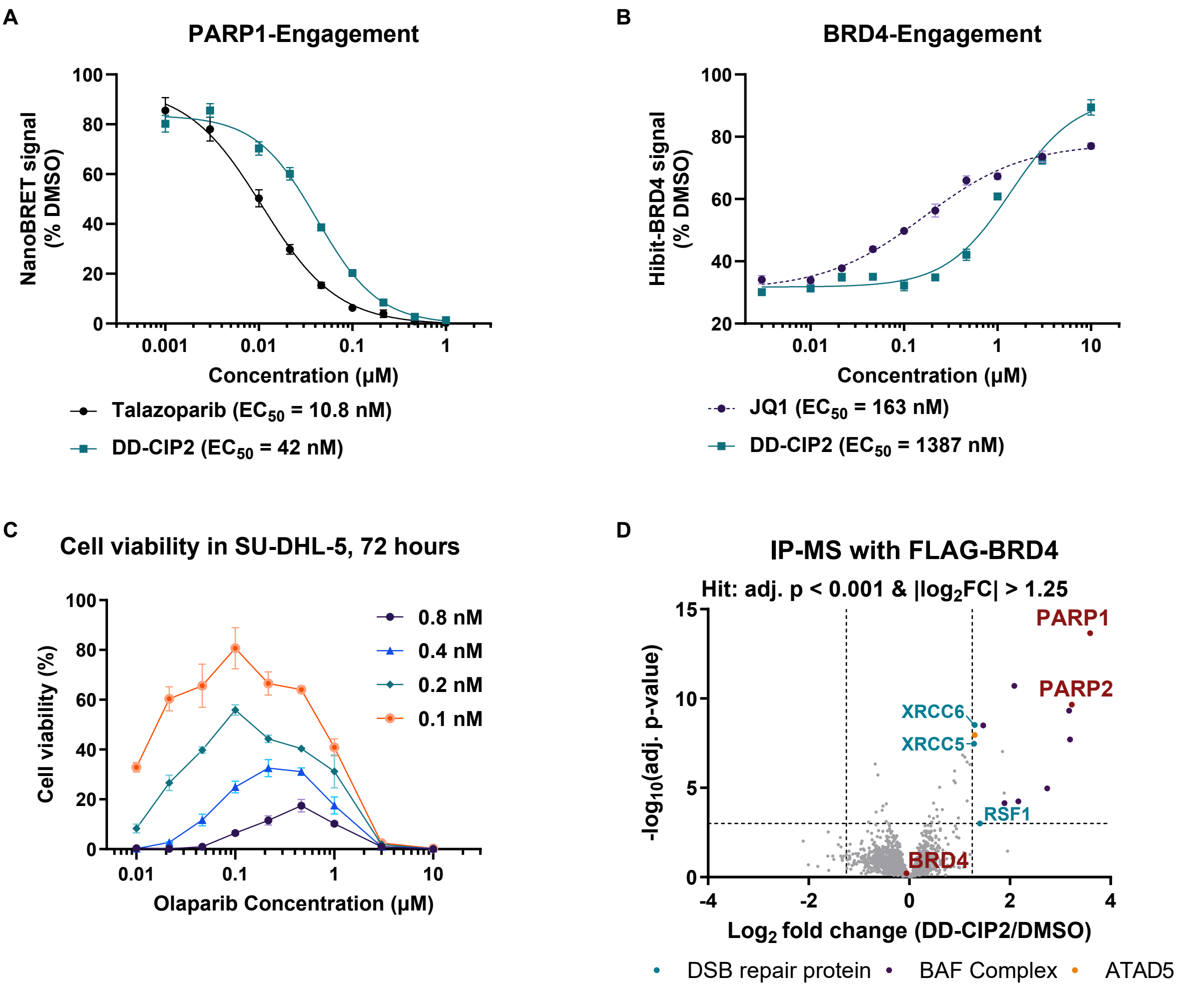


**Figure S6.** Characterization of DD-CIP2. A) PARP1 target engagement assay in HEK293T cells measured by Talazoparib or DD-CIP2 competitive displacement of Tracer-PARP-01 (NanoBRET signal normalized to DMSO, data are shown as means ± SEM, n=3 biological replicates). D) BRD4 target engagement assay in Hibit-BRD4 Jurkat cells measured by JQ1 or DD-CIP2 competitive displacement of dBET6 (HiBiT luminescence normalized to DMSO control with no dBET6, data are shown as mean ± SEM, n=3 biological replicates). Note data in Figure 1D is taken from the same experiment (JQ1 is repeated in both panels). C) Measurement of SU-DHL-5 cell viability after competitive titration of Olaparib to constant 0.1, 0.2, 0.4 or 0.8 nM DD-CIP2; cells were treated simultaneously with DD-CIP2 and Olaparib for 72 hours. Data are shown as means ± SEM; n = 3 biological replicates. D) FLAG IP-MS from HEK293T cells overexpressing FLAG-BRD4 treated with 100 nM DD-CIP2 for 1 hour; plotted with cut-offs of $|\log_2(\text{fold change})| \geq 1.25$ and adj. $P \leq 0.001$; 3 biological replicates. All P values were adjusted using Benjamini-Hochberg method from LIMMA-moderated t-test.

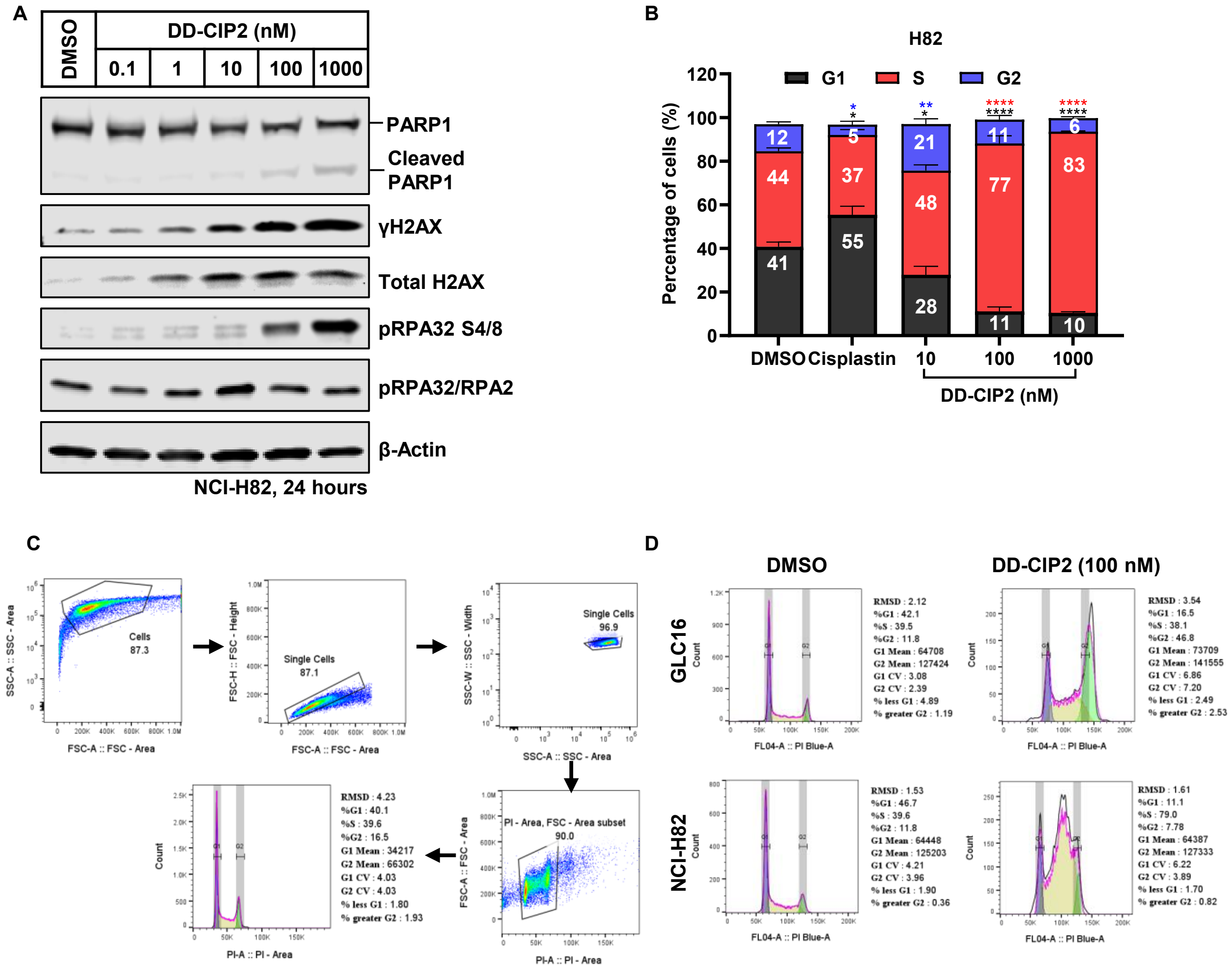


**Figure S7.** Characterization DD-CIP2 efficacy in SCLC cells. A) Western blots showing apoptosis (cleaved PARP1) and DNA damage signal (γH2AX, pRPA32 S4/8) markers in NCI-H82 cells treated with the indicated concentrations of DD-CIP2 for 24 hours. B) Cell cycle analysis of NCI-H82 cells treated with 10 μM Cisplatin or DD-CIP2 at indicated concentrations for 24 hours. Data are shown as means ± SEM, n = 3-5 biological replicates. Statistical analysis: *p < 0.05, **p < 0.01, ****p < 0.0001, one-way ANOVA with Bonferroni's post-test. C) Gating strategy for flow cytometry analysis of cell cycle. D) Representative histogram of cell cycle analysis for GLC16 and NCI-H82 cells treated with DMSO and 100 nM DD-CIP2.

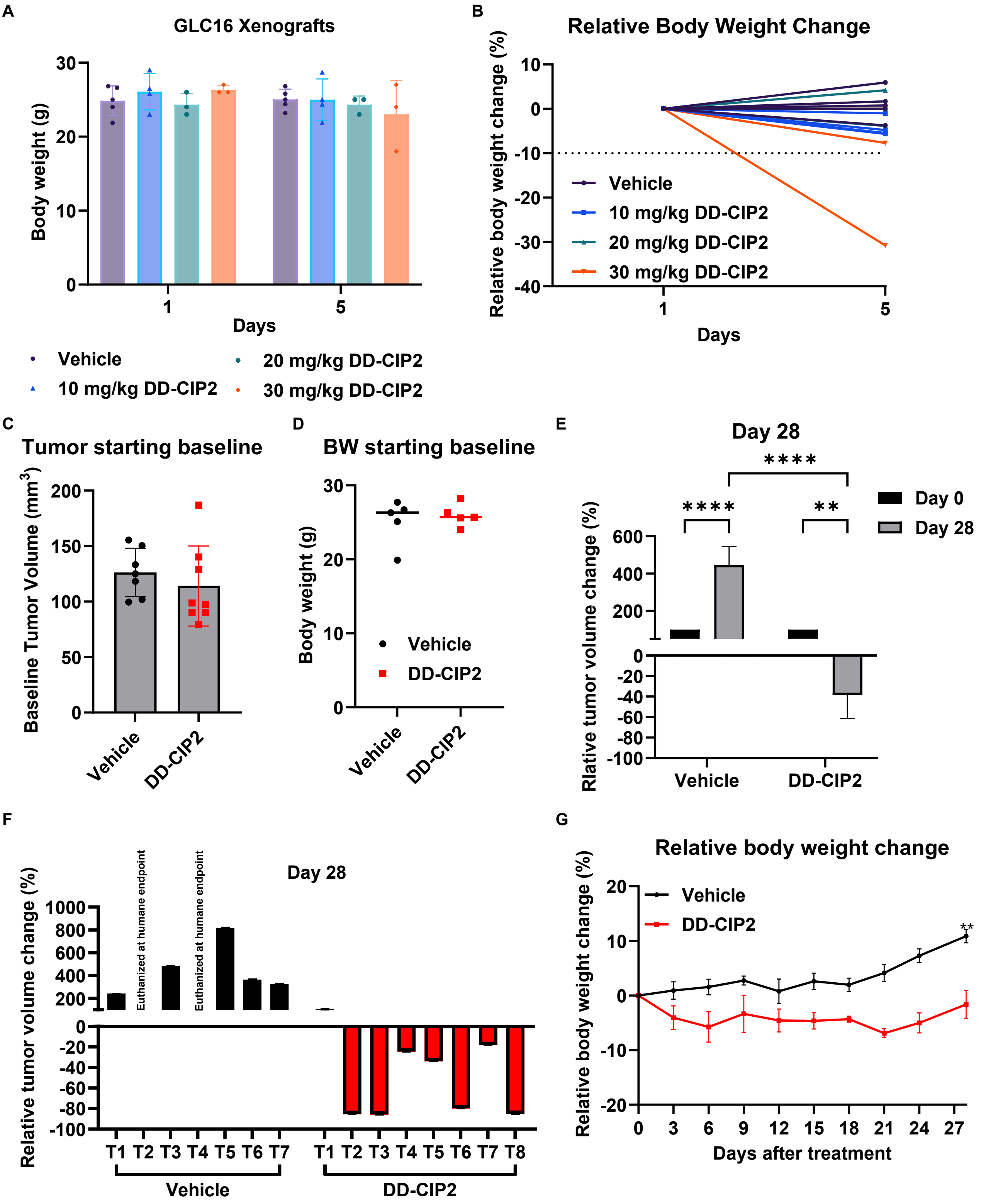
A
GLC16 Xenografts
Body weight (g)
Days
Vehicle
10 mg/kg DD-CIP2
20 mg/kg DD-CIP2
30 mg/kg DD-CIP2
B
Relative Body Weight Change
Relative body weight change (%)
Days
C
Tumor starting baseline
Baseline Tumor Volume (mm3)
Vehicle
DD-CIP2
D
BW starting baseline
Body weight (g)
E
Day 28
Day 0
Day 28
Rlative tumor volume change (%)
F
Day 28
Euthanized at humane endpoint
Relative tumor volume change (%)
T1 T2 T3 T4 T5 T6 T7
T1 T2 T3 T4 T5 T6 T7 T8
G
Relative body weight change
Relative body weight change (%)
Days after treatment

**Figure S8.** DD-CIP2 effectively induces DNA damage responses and anti-tumor efficacy in SCLC model *in vivo*. A) Mice body weight change at Day 1 and Day 5 in the PD study. Data represents means ± SEM, each dot represents individual mice. B) Spider plots for the percentage of body weight change at Day 1 and Day 5 in the PD study. C-D) The tumor volume (C) and the mice body weight (D) at the beginning of the DD-CIP2 treatment in the efficacy study. E) Relatively tumor volume change between Day 0 and Day 28. Data represents means ± SEM; n = 5-8 tumor samples per group; Statistical significance: $^{**}p < 0.01$, $^{****}p < 0.0001$, with Student's t-test. F) Relatively tumor volume change for individual tumors between Day 0 and Day 28. 10. G) Relative body weight change compared to day 0 shown as means ± SEM; n = 5 mice per group; Statistical significance: $^{**}p < 0.01$, with Student's t-test.
